# Copper drives prion protein phase separation and modulates aggregation

**DOI:** 10.1101/2023.02.15.528739

**Authors:** Mariana Juliani do Amaral, Aline Ribeiro Passos, Satabdee Mohapatra, Taiana Sousa Lopes da Silva, Renato Sampaio Carvalho, Marcius da Silva Almeida, Anderson de Sá Pinheiro, Susanne Wegmann, Yraima Cordeiro

## Abstract

Prion diseases are characterized by prion protein (PrP) transmissible aggregation and toxicity in the brain. The physiological function of PrP seems related to sequestering and internalization of redox-active Cu^2+^. It is unclear whether Cu^2+^ contributes to PrP aggregation, recently shown to be mediated by PrP condensation. We investigated the role of Cu^2+^ and oxidation in PrP condensation and aggregation using multiple biophysical and biochemical methods. We find that Cu^2+^ promotes PrP condensation at the cell surface and *in vitro* through co-partitioning. Molecularly, Cu^2+^ inhibited PrP β-structure and hydrophobic residues exposure. Oxidation, induced by H_2_O_2_, triggered liquid-to-solid transition of PrP:Cu^2+^ condensates and promoted amyloid-like PrP aggregation. In cells, overexpression of PrP^C^ initially protected against Cu^2+^ cytotoxicity but led to PrP^C^ aggregation upon extended copper exposure. Our data suggest that PrP condensates function as a buffer for copper that prevent copper toxicity but can transition into PrP aggregation at prolonged oxidative stress.

## Introduction

The prion protein (PrP) is highly expressed in neurons of the central nervous system. PrP is an extracellularly presented GPI-anchored plasma membrane (PM) protein that is enriched at pre-synaptic sites, and about 10-20% of total PrP is found in the cytosol^1–3^. In prion diseases, PrP undergoes self-propagating misfolding into a pathogenic β-sheet enriched form that is associated with scrapie prion disease (PrP^Sc^). Prion diseases are fatal, and the only known transmissible proteinaceous diseases in humans and other mammals^1^.

PrP^Sc^-infected cell models and mice show metal ion (e.g. Cu^2+^, Zn^2+^, Fe^2+^, Mn^2+^) dyshomeostasis and reduced cellular response to oxidative stress^4–6^. Moreover, the lack of PrP^C^ in *prnp* neurons from knockout mice leads to enhanced susceptibility to oxidative stress^7,8^. Copper binding to PrP was described *in vitro^9^* and *in vivo^10^* and occurs with different affinities depending on Cu^2+^ availability, and exhibits negative cooperativity^11,12^, which suggests a role of PrP as a Cu^2+^ buffering system^11,13^ and in other biological functions^14^. For example, PrP appears to facilitate the internalization of Cu^2+^, whereby a reduction to Cu^+^ occurs in acidic endosomes^9,12^. Furthermore, PrP has been suggested to ‘quench’ redox activity of Cu^2+^ and other neurologically relevant transition metals (Fe, Zn, Mn)^12,15,16^.

Nine solvent-exposed histidine residues in PrP are involved in Cu^2+^ coordination. The main Cu^2+^ binding activity is encoded in the PrP N-terminal domain whereby 4 histidine residues (H61, H69, H77, H85; human protein residue numbering) within the octarepeat region (60-91; 4 PHGGGWGQ tandem repeats) and two nearby histidines (H96 and H111) coordinate copper ions (at pH~7)^12^. Recently, two C-terminal histidines, H140 and H177, were shown to co-bind Cu^2+^, thereby tethering PrP N- and C-terminal domains and promoting their interaction^17^. Additionally, H187 was also shown to coordinate Cu^2+^ ^18^. The Cu^2+^ coordinating octarepeat region of PrP was predicted to contain a distinctive uniform distribution of aromatic residues^19^ and to form low-complexity amyloid-like kinked segments (LARKS)^20^, which establish transient β-sheet interactions and seem to be involved in driving liquid-liquid phase separation (LLPS) of low complexity domains (LCDs) in proteins forming biomolecular condensates, *e.g.*, membrane-less organelles. Protein condensates are dynamic liquid-like assemblies that can act as reaction hubs or structural entities, and contribute to the spatiotemporal organization of cellular processes^21,22^. However, for proteins related in different neurodegenerative diseases, e.g., fused in sarcoma (FUS) and hnRNPA1 in amyotrophic lateral sclerosis (ALS), TDP-43 in frontotemporal dementia and ALS, α-synuclein in Parkinson’s disease, and Tau in Alzheimer’s disease, a transition from liquid-like condensates to gel- or solid-like structures has been described and linked to their aggregation and neuropathological potential^23,24^. PrP was shown to undergo LLPS *in vitro*, modulated by nucleic acids, other proteins, and the physicochemical environment^25,26^.

We hypothesized that Cu^2+^, via its multivalent interactions with LARKS-containing octapeptide region and other domains of PrP, could promote physiological PrP condensation, and even be involved in driving pathological PrP aggregation. Using a multiparametric biophysical and biochemical approach combining *in vitro* and live cell data, we, indeed, revealed that Cu^2+^ promotes the condensation of recombinant PrP (PrP) *in vitro*, as well as membrane anchored PrP^C^ in cells. *In vitro*, Cu^2+^ together with hydrogen peroxide, mimicking oxidative stress, decreased the molecular diffusion inside PrP condensates, followed by a liquid-to-solid transition and the formation of amyloid-like PrP aggregates enriched in di-tyrosine crosslinks. In cells overexpressing fluorescently labeled PrP^C^, Cu^2+^ promoted the formation of membrane-bound and cytoplasmic PrP^C^ condensates. Interestingly, PrP^C^ overexpression protected against Cu^2+^ cytotoxicity, indicating that PrP^C^ condensation may function as copper buffering system by sequestering Cu^2+^ ions. Our data indicate that PrP condensation may be an integral mechanism for a variety of copper- and other metal ion-related functions, such as metal ion uptake, transport, storage, and gradual release, which are important processes in signaling and oxidation protection. However, ROS production due to prolonged Cu^2+^ or H_2_O_2_ exposure can induce a liquid-to-solid transition or PrP:Cu^2+^ condensates, which can drive pathological PrP aggregation.

## Results

### PrP^C^ undergoes membrane associated condensation in cells

PrP^C^ is an extracellular PM protein that carries a glycosylphosphatidylinositol anchor (GPI)-anchor on its C-terminus, which is essential for the intracellular trafficking of PrP^C^ from the ER through the Golgi, and its anchoring to the PM^27^. Whether and where - at the membrane or in the cytosol - PrP^C^ forms condensates in cells is not known. To assess PrP^C^ phase separation, we used a construct containing PrP^C^ fused to YFP followed by a GPI-anchor (PrP^C^-YFP-GPI) and expressed it in HEK293 cells.

Live cell confocal microscopy showed that most of PrP^C^-YFP-GPI localized to the PM (**Fig. 1a**, top left). Treatment with 300 μM CuCl_2_ (from here on: Cu^2+^) for 1 hour appeared to increase the accumulation of PrP^C^-YFP-GPI at the PM, especially pronounced at cell-cell interface, where the PMs of two adjacent cells expressing PrP^C^-YFP-GPI were in contact (**Fig. 1a**, bottom left). Line plots across cell-cell interfaces and individual PMs of the same cells (examples in **Fig. 1b**) revealed a ~2-fold higher PrP^C^-YFP-GPI fluorescence intensity at cell-cell interfaces (II) compared to the sum of the respective individual membranes (I_M1_+I_M2_), indicating a cooperative assembly of PrP^C^-YFP-GPI at the interface of Cu^2+^-treated cells (simple addition of PrP^C^-YFP-GPI signal from adjacent PMs would give values of I_I_/(I_M1+_I_M2_) ≤ 1) (**Fig. 1c**). In untreated cells, the cell-cell interface had only 1.2-fold higher intensity indicating a less cooperative PrP^C^-YFP-GPI assembly (I_J_/(I_M1+_I_M2_) for no Cu^2+^ vs. Cu^2+^: P= 0.0001). The PrP^C^-YFP-GPI expression levels and absolute fluorescence intensity were similar between treated and untreated cells (**Extended data Fig. 1c**). Cells expressing YFP-GPI did not show enrichment at the cell-cell interface (**Fig. 1c**).

**Fig. 1.**
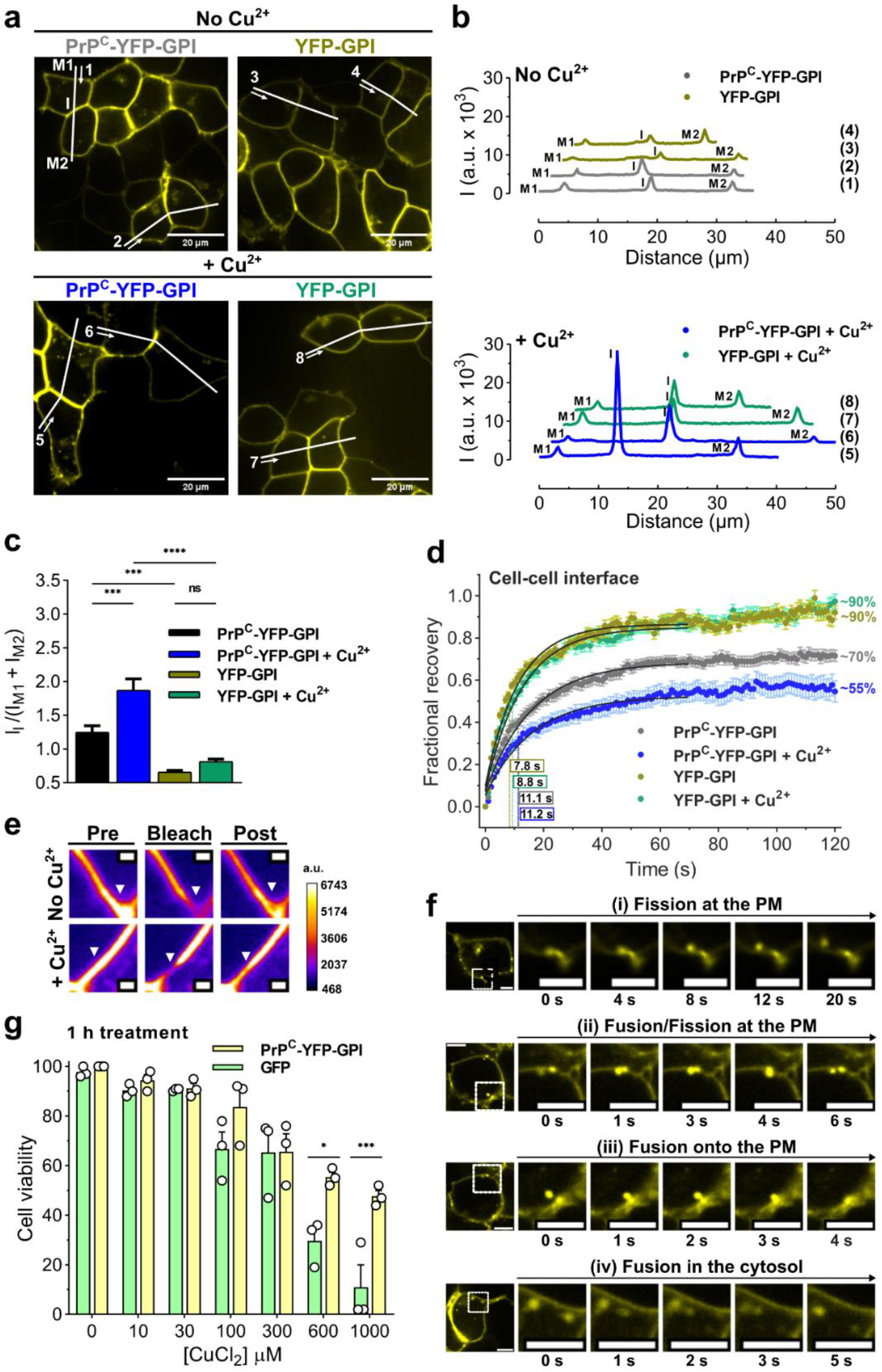
PrP^C^ condensation at cell-cell contacts is promoted by Cu^2+^. **a**, Representative confocal images showing HEK293 cells transfected with PrP^C^-YFP-GPI or YFP-GPI (for 48 h) show localization at the PM, and accumulation of PrP^C^-YFP-GPI at cell-cell interfaces in untreated (top) Cu^2+^-treated (300 μM CuCl_2_) cells (bottom). White lines across cell membranes (M1, M2) and interfaces (I) indicate positions and numbering of line plots in ‘b’. Arrows mark the start and direction of measurement. **b**, Line plots from ‘a’ showing PrP^C^-YFP-GPI or YFP-GPI fluorescence intensity of individual PMs (M1, M2; not in contact with other cells) and the cell interface (I) of two adjacent cells treated (+ Cu^2+^) or not (no Cu^2+^). **c**, Quantification of fluorescence intensity at the cell-cell interface relative to the sum of intensities measured for individual PMs of the contacting cells (I_I_/(I_M1+_I_M2_)). Data shown as mean ± SEM, n=21 (PrP^C^-YFP-GPI), n=20 (PrP^C^-YFP-GPI + Cu^2+^), n= 23 (YFP-GPI) and n=20 (YFP-GPI + Cu^2+^). One-way ANOVA with Tukey test. ****P < 0.0001; ***P<0.001; ns, not significant **d**, FRAP shows a lower recovery of PrP^C^-YFP-GPI at the cell-cell interface of Cu^2+^-treated cells, indicating less molecular diffusion upon Cu^2+^-treatment, as opposed to YFP-GPI with Cu^2+^ treatment or not. n = 26 cell-cell interfaces per group. Fitting to obtain times to half recovery (t1/2; values inside rectangles) are shown in black. **e**, Example of circular areas of PrP^C^-YFP-GPI in the cell-cell junctions (pre, left) that have been bleached (bleach, middle) and their recovery followed by 120 seconds (post, right). Pseudocolored intensity scale shown on the right. **f**, Representative time-lapse confocal image series of Cu^2+^-treated cells showing liquid-like fission (i) and fusion (ii and iii) of small PrP^C^-YFP-GPI clusters at the PM, and in the cytosol (iv). Imaging time points indicated below. **g**, Cell viability assay of HEK293 cells transfected with PrP^C^-YFP-GPI or GFP following treatment with increasing concentrations of CuCl_2_ for 1 h. *P<0.05, ***P< 0.001. Data shown as mean ± SEM, n=3 independent replicates, one-way ANOVA (Sidak). Scales bars are 20 μm in (a), 1 μm in (e) and 5 μm (f).

Assembly cooperativity is a characteristic of condensation, whereby proteins concentrate in a dense liquid-like phase. Proteins inside biomolecular condensates show a reduced molecular diffusion compared to diffuse proteins in the cytosol, which can be observed as slower and/or less fluorescence recovery after photobleaching (FRAP). Condensates of membrane proteins show an additional FRAP delay because of restricted 2D-diffusion^28^. To test whether Cu^2+^-promoted enrichment of PrP^C^-YFP-GPI at cell-cell interfaces may be driven by condensation of extracellularly presented PrP^C^, we compared FRAP of PrP^C^-YFP-GPI in untreated and Cu^2+^-treated cells (**Fig. 1d**). In response to Cu^2+^, PrP^C^-YFP-GPI showed significantly less recovery (no Cu^2+^: ~70% vs. Cu^2+^: ~55%; P=0.0054) indicating less mobile PrP^C^ species inside assemblies. FRAP of YFP-GPI was as expected for non-condensating membrane proteins and did not change by Cu^2+^ treatment (**Fig. 1d**). In addition, confocal time lapse imaging revealed both fusion and fission of small PrP^C^-YFP-GPI clusters in the PM, and Cu^2+^ promoted puncta in the PM (**Fig 1f** and **Extended Data Fig. 1d**). Importantly, compared to GFP, PrP^C^-YFP-GPI expressing cells showed a higher viability at increasing CuCl_2_ concentration in the culture medium (**Fig. 1g** and **Supplementary Fig. 1;** P=0.0111 at 600 μM and P=0.002 at 1 mM CuCl_2_), indicating that PrP^C^ exerted a protective effect against Cu^2+^-induced cytotoxicity.

In addition to the PM-bound main pool, we observed a small fraction of PrP^C^-YFP-GPI as highly mobile, small (diameter <1μm), round structures in the cytosol, often near to the PM (**Extended Data Fig. 1b**; **Supplementary Video 1**), which is consistent with previously reported cytosolic PrP^C^ ^29^. Time lapse imaging showed coalescence of cytosolic PrP^C^-YFP-GPI and the exchange of PrP^C^-YFP-GPI signal between cytosolic structures and the PM within seconds - in both directions (**Fig. f**; **Extended Data Fig. 1d**; **Supplementary Videos 1** and **2**). These events were reminiscent of condensate fusion and fission. In some cells, we observed large accumulation of PrP^C^ localized next to the nucleus (**Extended Data Fig. 1b**) corroborant to previous observations on PrP overexpression^30^, which may be related to misfolded protein accumulation at the aggresome^2^. Of note, cells maintained in RPMI 1640 (containing 10% FBS), Opti-MEM or Imaging solution showed similar subcellular distribution of PrP^C^-YFP-GPI (**Supplementary Fig. 1c**).

Together, our data indicate that the presence of Cu^2+^ enhances the accumulation of PrP^C^ at cell-cell interfaces driven by condensation, and, simultaneously, reduces PrP^C^ molecular diffusion inside condensates. Sequestration of Cu^2+^ into PrP^C^ condensates could reduce oxidative stress, which would explain the increased cell viability upon copper treatment in the presence of PrP^C^.

### Cu^2+^ triggers recombinant PrP condensation at physiological conditions in vitro

We proceeded to investigate the molecular mechanisms of Cu^2+^-driven PrP condensation *in vitro*. Different bioinformatic prediction tools for the LLPS propensity of proteins (Fuzdrop^31^, catGRANULE^32^, ParSe^33^, and PScore^34^) agreed that the octarepeat region of PrP (residues 60-91), which binds Cu^2+^ ions via 4 histidine residues, has the highest probability to undergo LLPS (**Fig. 2a**), probably because of the five LARKS. Interestingly, LLPS predictions for PrP mutants with extra octarepeats that cause early-onset inherited prion diseases^35^ show enhanced LLPS propensities (**Extended data Fig. 2a**), highlighting the relevance of this region for both LLPS and pathology and suggesting a modulation of PrP phase transitions by copper.

**Fig. 2.**
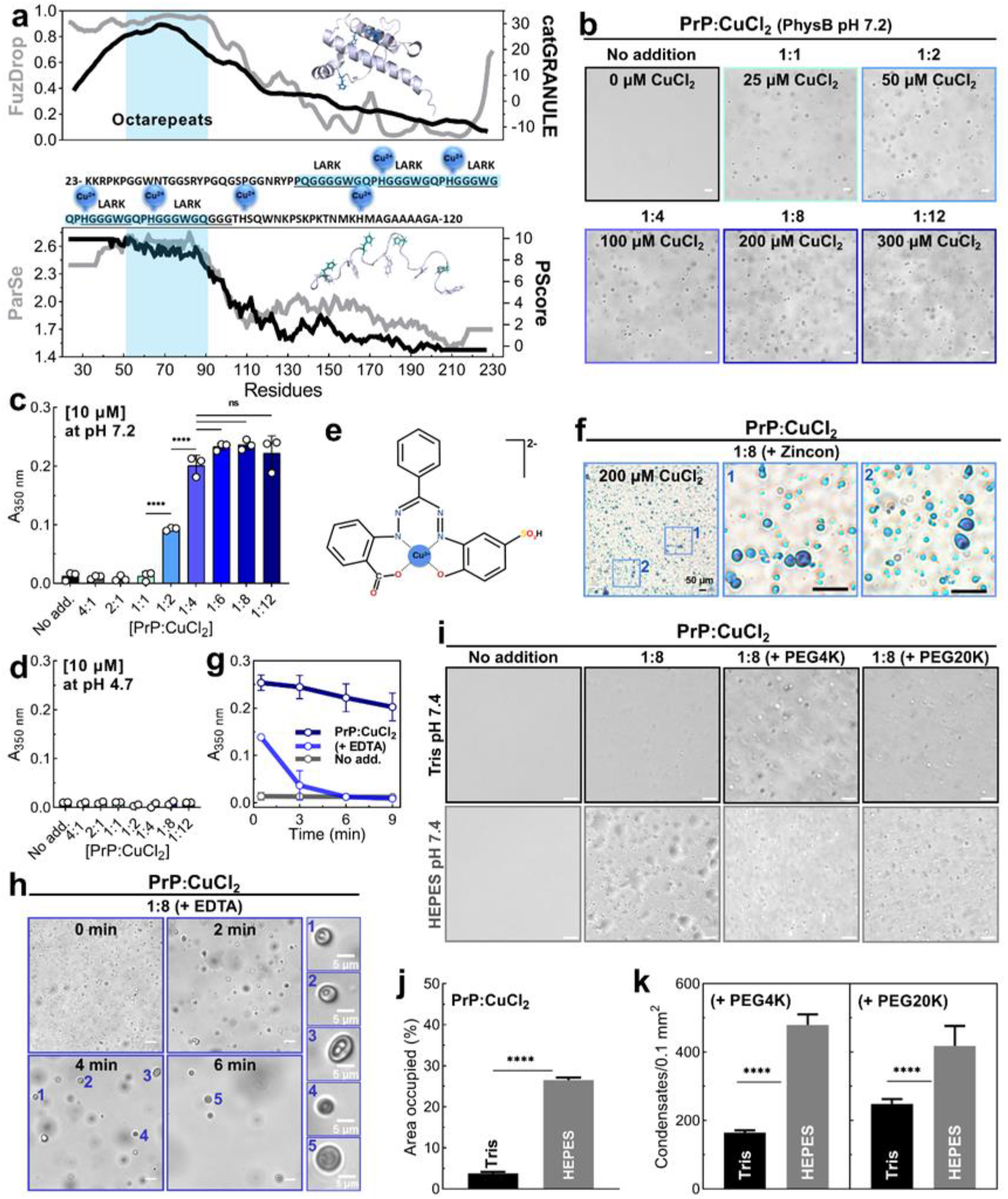
Cu^2+^ drives recombinant PrP condensation. **a**, LLPS prediction of PrP (UniProt ID P04156) by FuzDrop^31^ (top, gray line, left axis), catGRANULE^32^ (top, black line, right axis), ParSe^33^ (bottom, gray line, left axis) and PScore^34^ (bottom, black line, right axis). Middle: Amino acid sequence of PrP N-terminal domain (23-120) with indicated copper-coordinating histidine residues. Five octarepeats (highlighted in blue) are predicted to be low-complexity amyloid-like kinked segments (LARKS) forming transient contacts involved in LLPS. Top inset: PrP C-terminal domain structure (residues 121-231, PDB 1XYX) with Cu^2+^-coordinating His side chains in blue. Bottom inset: octarepeats structure (residues 23-106, PDB 2KKG) with evenly interspaced Trp (green) and Tyr (white) side chains that potentially contribute to LLPS. **b**, Phase contrast images of PrP at 25 μM incubated for 2 h with 25-200 μM CuCl_2_. **c**, **d**, Turbidity (absorption at 350 nm) of 10 μM PrP at increasing CuCl_2_ concentration at pH 7.2 (c) and pH 4.7 (d). **e**, Structure of Zincon complexed to Cu^2+^. **f**, Light microscopy images of 25 μM PrP with 200 μM CuCl_2_ captured by a color high-resolution camera after 30 min preparation followed by addition of Zincon. Middle and left images show close-up of marked regions. **g**, Turbidity kinetics (absorbance at 350 nm) of 10 μM PrP only, with 120 μM CuCl_2_ or upon addition of 1.2 mM EDTA to PrP:CuCl_2_. **h**, Representative images from time-lapse of 25 μM PrP with 200 μM CuCl_2_ (1:8 molar ratio) upon EDTA addition (2 mM; 10 × molar excess in relation to Cu^2+^). Right: close-up showing vacuolated condensates. **i,** 25 μM PrP with 200 μM CuCl_2_ in 25 mM of specified buffers (without salt) containing no, or 10% PEG4K or PEG20K. Tris-buffer shows less condensates than HEPES, likely because Tris coordinates Cu^2+^. Samples prepared at 25°C for 1 h before imaging. **j**, Quantification of the surface area occupied by PrP:CuCl_2_ condensates shown in (i) (second column). **k**, Quantification of condensates’ number shown in (i) (third and fourth columns). Data in (c, d, g) shown as mean ± SD, n= 3 replicates, 2 in (d) and 6 images in (j, k). All experiments were performed at 25°C in PhysB unless otherwise stated. Data analyzed by one-way ANOVA with Tukey test (c) or two-tailed Student’s t-test (j, k). ****P < 0.0001; ns, not significant. Scales bars are 10 μm in (b, h, i) and 50 μm in (f).

To examine how the binding of Cu^2+^ - the best-known function of the octarepeat region^10,12^-would modulate PrP condensation, we incubated recombinant PrP in physiological ion and molecular crowding conditions (PhysB; incl. 6% PEG4K; 25°C; **Supplementary Table 1**) with different concentrations of CuCl_2_ and monitored LLPS by microscopy (25 μM PrP; **Fig. 2b**) and turbidity (10 μM PrP; **Fig. 2c**). Whereas PrP alone did not phase separate in these conditions, the addition of equimolar or higher amounts of CuCl_2_ triggered the rapid formation of PrP condensates. PrP turbidity measurements indicated that LLPS plateaued at a PrP:CuCl_2_ molar ratio of 1:4. Tryptophan fluorescence anisotropy (mouse PrP has eight tryptophan residues, being seven in the N-terminal region) confirmed the formation of larger species (e.g., oligomers or condensates) at PrP:CuCl_2_ molar ratios of 1:4 and 1:8 (**Extended data Fig. 2a**). At pH 4.7, when the octapeptide histidine residues are protonated and metal coordination is negligible^36,37^, no condensation occurred (pKa of octarepeat histidines= 6-7; pKa of H96 and H111=7-8) (**Fig. 2d**), even with higher molecular crowding (10% PEG4K or PEG20K) (**Supplementary Fig. 2b**) or low temperature (**Supplementary Fig. 2c**). Of note, at non-physiological conditions of high protein concentrations (~70 μM at ~13°C; **Supplementary Fig. 2e**), very high molecular crowding (10% PEG20K; **Extended data Fig. 2d, e**), and low temperatures (25 μM at ~10°C; **Supplementary Fig. 2c**), we were able to observe ‘homotypic’ condensation for PrP alone, which seemed to be inhibited at low pH 4.7 (**Supplementary Fig. 2b, c**) and promoted by high NaCl (300-500 mM) in the buffer (**Supplementary Fig. 2e**).

To ascertain whether Cu^2+^ ions are partitioning into condensates, we used the dye Zincon, which forms a blue complex when bound to copper^38^ (**Fig. 2e**). Microscopy (**Fig. 2f**) and spectrophotometric quantification of the dense phase with Zincon revealed that Cu^2+^ ions enriched at least 10-fold in the PrP:CuCl_2_ condensates (**Supplementary Fig. 3**). To test whether the Cu^2+^ in the condensates was essential for their stability, we added ethylenediaminetetraacetic acid (EDTA), a chelator with high affinity to Cu^2+^, to preformed PrP:Cu^2+^ condensates, expecting that EDTA would dissolve the condensates by complexing Cu^2+^ necessary for condensation. Indeed, turbidity measurements suggested that EDTA dissolved PrP:Cu^2+^ condensates within a few minutes (**Fig. 2g**). Microscopy showed the formation of larger vacuolated intermediates that gradually dissolved (**Fig. 2h**). Likewise EDTA, Tris coordinates Cu^2+^ ions^39^ and has been shown to outcompete Cu^2+^-binding to octarepeat peptides derived from PrP^40^. We indeed found that Tris could interfere with PrP:CuCl_2_ LLPS (1:8 molar ratio) (**Fig. 2i**). Comparing to ‘Cu^2+^ non-coordinating’ HEPES buffer^41^, 6-times less PrP:CuCl_2_ condensates were observed in Tris-buffer (**Fig. 2j, k**).

We further investigated the conditions at which CuCl_2_-induced PrP condensation would occur. At a fixed, physiological relevant concentration of CuCl_2_ (80 μM; **Supplementary Table 1**), LLPS depended on PrP concentration and started at 4 μM PrP (**Extended data Fig. 2b**). Next, we used a bis-glycine complex (Cu(Gly)_2_) as source of Cu^2+^ ions to resemble the cellular milieu, in which Cu^2+^ is mostly found protein bound. Even in the Cu(Gly)_2_ form, Cu^2+^ was able to induce PrP LLPS (**Extended data Fig. 2c**). In the same experiment we also observed the dependence of PrP:Cu(Gly)_2_ LLPS on increasing molecular crowding (% PEG4K), indicating a synergy between Cu^2+^ and molecular crowding in driving PrP LLPS (**Extended data Fig. 2c**). The amount of molecular crowding induced by PEG with different molecular weights (**Extended Data Fig. 2d-g**) appeared to influence the surface attachment and shape of PrP:Cu^2+^ condensates.

Altogether, these data show that Cu^2+^ is a major modulator of PrP phase separation, which confirms our observations made for PrP^C^ in cells. PrP:Cu^2+^ condensation occurs at physiological PrP, copper, pH, salt, and crowding conditions.

### Physicochemical properties of PrP:CuCl_2_ condensation

We sought to understand which molecular interactions would drive and stabilize PrP:CuCl_2_ condensates. FRAP suggested liquid-like molecular diffusion of PrP inside condensates (**Fig. 3a**). Interestingly, turbidity temperature ramping experiments (between 13 and 40°C) showed that Cu^2+^-induced PrP condensation increased with decreasing temperatures, starting at ~24°C, which was fully reversible upon re-heating the sample (**Fig. 3b**). Dynamic light scattering (DLS) confirmed that the majority of PrP:CuCl_2_ condensates were larger at 20°C (hydrodynamic radius, Rh~540 nm) than at 37°C (Rh~340 nm) (**Fig. 3c**), corroborating the induction of PrP condensation at lower temperatures (**Fig. 3b**). Of note, in PhysB in the presence of molecular crowding (PhysB with 6% PEG4K), a small amount (mass percentage <1%) of high-order oligomers occurred (**Fig. 3h**).

**Fig. 3.**
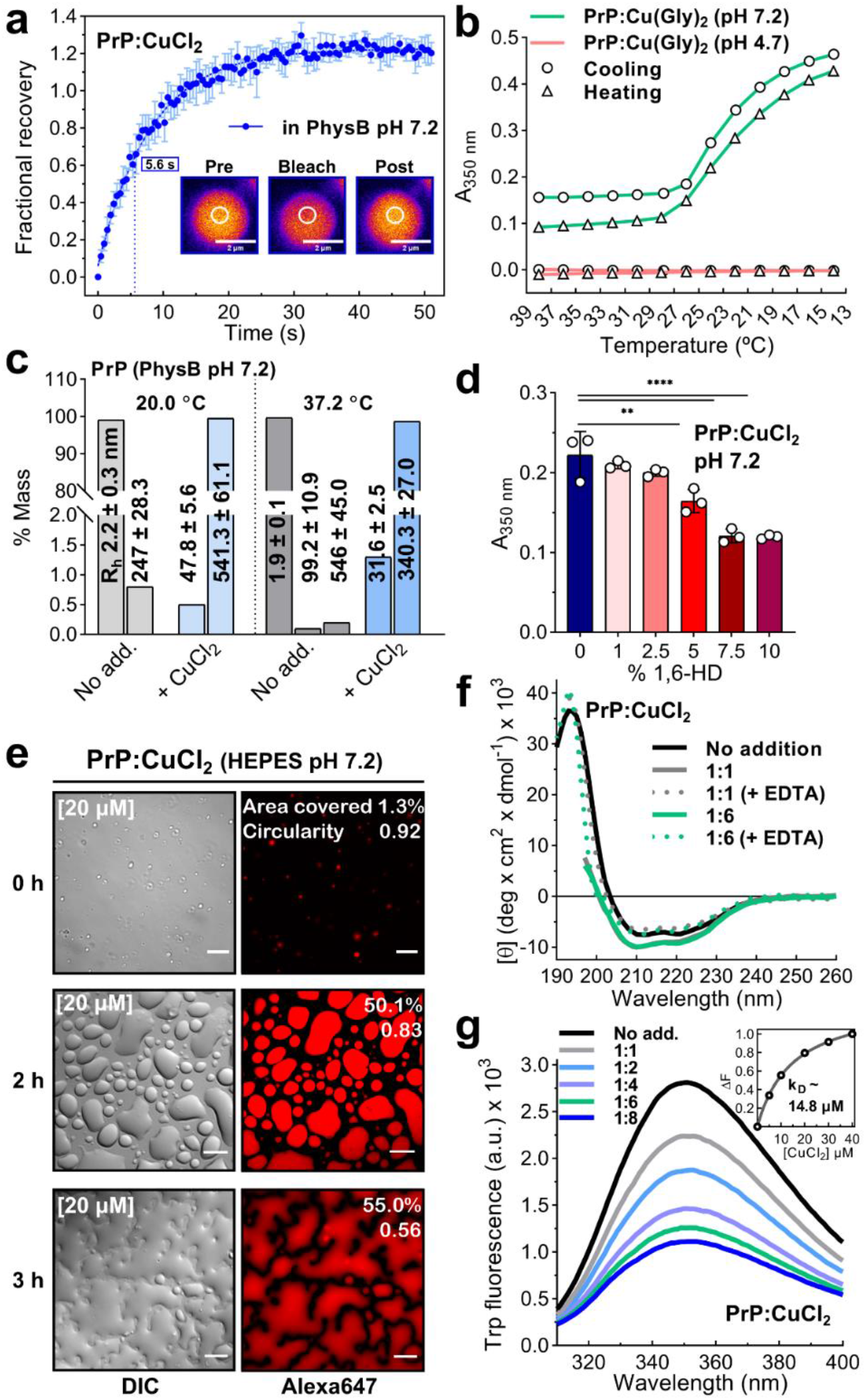
Physicochemical characterization of Cu^2+^-driven PrP condensation. **a**, FRAP of 10 μM PrP (spiked with 0.1% PrP-Alexa647) with 80 μM CuCl_2_ (1:8 molar ratio), performed immediately after sample preparation. Data shown as mean ± SEM (n=3 condensates). Time to half recovery (t1/2) is shown in the rectangle. **b**, Turbidity of 25 μM PrP with 150 μM Cu(Gly)_2_ (1:6) in PhysB prepared with HEPES (pH 7.2, green) or MES (pH 4.7, red) as a function of decreasing (circle, cooling) or increasing (triangle, heating) temperature. **c,** DLS of 25 μM PrP only (grey) or with 200 μM CuCl_2_ (blue; 1:8 molar ratio) at 20°C and 37.2°C. Data show mass percentage of each population having the indicated R_h_ (values in nm inside/above bars). **d**, Turbidity of PrP:CuCl_2_ (1:8 molar ratio) condensation as a function of increasing 1,6-hexanediol (1,6-HD) concentrations (measured after 6 min). Data shown as mean ± SD (n=3). One-way ANOVA (Tukey). ****P < 0.0001, **P < 0.01. **e**, DIC and fluorescence images of 20 μM PrP (spiked with 0.1% PrP-Alexa647) with 160 μM CuCl_2_ (1:8 molar ratio) show wetting of poly-D-lysine coated glass surfaces over time. **f**, Far-UV Circular Dichroism spectra of unbound 2.5 μM PrP (black) alone or with 2.5 μM CuCl_2_ (gray), and upon addition of EDTA (7.5 μM; dotted gray); with 15 μM CuCl_2_ (green), and upon addition of EDTA (45 μM; dotted green). **g**, Quenching of tryptophan fluorescence (excitation at 295 nm; emission 310-400 nm) of 5 μM PrP with 5-40 μM CuCl_2_. **inset**, changes in fluorescence intensity area (ΔF) as a function of CuCl_2_ concentration. Dissociation equilibrium constant (kD) determination from hyperbolic fitting. All spectra acquired at 25°C in PhysB, unless otherwise specified. Scales bars, 2 μm (a), 10 μm (e).

Protein condensation is driven by weak multivalent intra- and intermolecular interactions between residue side chains or protein structural elements^29^. Clients of protein condensates, such as other proteins, nucleic acids, or ions, can modulate condensate stability by promoting or disturbing these interactions. To test whether hydrophobic interactions stabilize PrP:CuCl_2_ condensates, we added the aliphatic alcohol 1,6-hexanediol (1,6-HD) to preformed PrP:CuCl_2_ condensates. 1,6-HD was reported to dissolve condensates of PrP^90-231^ (residues 90 to 231) at 10% (w/v)^26^, and of PrPY145Stop (residues 23 to 144)^42^ at 2% (w/v), highlighting the role of weak hydrophobic contacts in homotypic LLPS of these constructs. For Cu^2+^-driven PrP LLPS, we observed only 45% decrease in turbidity at 10% 1,6-HD and no effect at 2% 1,6-HD (**Fig. 3d**). We concluded that hydrophobic interactions have a smaller contribution to PrP:CuCl_2_ LLPS, and electrostatic interactions may play an important role. Indeed, in the absence of counter ions in the buffer that weaken electrostatic interactions (10 mM HEPES, pH 7.2, no salt), Alexa647-PrP:CuCl_2_ condensation was enhanced and pronounced wetting of positively-charged (poly-D-lysine coated) glass surfaces occurred (**Fig. 3e** and **Supplementary Video 3**). We previously observed a similar behavior for electrostatically driven coacervates of Tau^43^.

Copper binding to the octarepeat domain of PrP may induce LARKS, which are transient kinked β-structures, and thereby promote PrP LLPS^44^. To examine whether Cu^2+^ induces conformational changes in PrP, we performed far-UV circular dichroism spectroscopy (**Fig. 3f**). The experiments were performed at PrP concentrations (2.5 μM) below the critical concentration for LLPS to avoid Cu^2+^-driven condensation, which could produce misleading conclusions. The α-helical content of PrP is characterized by negative bands at 222 and 208 nm, and a positive maximum at ~195 nm (**Fig. 3f**). Spectra deconvolution analysis by the Contin-LL software (dataset SMP180)^45^ estimated 33.6% of α-helical content, 27.4% β-sheets, 8.6% turns and 30.4% disordered structure, consistent with recombinant PrP CD spectrum^46^ and NMR-determined 3D structures (*e.g*., PDBs: 1QM0, 1HJM, 2LSB, 5YJ5). Upon addition of CuCl_2_ at 1:1 and 1:6 molar ratios, a minor increase of the 222 and 208-nm bands indicated further stabilization of α-helical segments and/or gain of other conformational elements (**Fig. 3f**). Intrinsic tryptophan fluorescence supported this idea by showing fluorescence quenching of PrP in the presence of CuCl_2_ (**Fig. 3g**), indicative of Cu^2+^-induced structural changes near tryptophan residues. The binding affinity of Cu^2+^ and PrP in PhysB (**Fig. 3g, inset**) supported previous results (*i.e*. K_D_ ~ 3-12 μM)^9,11,47^. These data corroborate the idea that Cu^2+^ may stabilize certain PrP conformations, probably through long-range interactions between the N- and the C-terminal domains^13,17^.

### Hydrogen peroxide and divalent transition metal ions modulate PrP phase transitions

Given that PrP^C^ binds other divalent metal ions and regulates their homeostasis^48^, we investigated how FeCl_2_, ferric ammonium citrate (FAC), ZnCl_2_, CoCl_2_, CdCl_2_ and MnCl_2_ would modulate PrP condensation. Phase contrast microscopy and turbidity measurements showed that FeCl_2_, FAC, ZnCl_2_, CdCl_2_ and MnCl_2_ trigger PrP LLPS (**Extended Data Fig. 3a-c**). Copper sulphate (CuSO_4_) promoted PrP LLPS like CuCl_2_, hence, Cu^2+^ effect on phase separation is not dependent on the type of salt counter anion (**Extended Data Fig. 3a-c**). ZnCl_2_ only triggered PrP phase separation at higher molar ratios (1:8) and not at equimolarity like CuCl_2_ or CuSO_4_, FeCl_2_, FAC, CdCl_2_ and MnCl_2_. Interestingly, CoCl_2_ did not trigger PrP LLPS at the conditions tested. It is worth noting that Cu^2+^, Fe^2+^, Zn^2+^, Co^2+^ and Mn^2+^ are essential trace elements in the nervous system, especially in the synaptic cleft. We speculate that synaptic PrP condensates could act as a buffer for regulating the concentration of divalent metal ions at synapses.

Cu^2+^ is redox active and can engage with hydrogen peroxide (H_2_O_2_) in Fenton reaction, generating reactive oxygen species (ROS). Since PrP:Cu^2+^ complexes can encounter H_2_O_2_ intra- and extracellularly to generate ROS, we investigated the effect of H_2_O_2_ on PrP LLPS in the presence or absence of Cu^2+^. Adding 10 mM H_2_O_2_ to PrP:Cu^2+^ LLPS conditions produced a mix of condensates and branched aggregates at all PrP:Cu^2+^ molar ratios after 2 h, most pronounced at higher copper concentrations (**Fig. 4a,b** and **Extended Data Fig. 4a,b**). H_2_O_2_ alone did not trigger PrP condensation or aggregation (**Supplementary Fig. 4b**). Condensates adhered to the branches of aggregates were visualized (**Fig. 4c**). Comparing turbidity measurements of PrP:Cu^2+^ LLPS, indicated that H_2_O_2_ significantly stimulated PrP:Cu^2+^ LLPS at 1:1 and 1:2 molar ratio PrP:Cu^2+^ compared to the absence of H_2_O_2_ (PrP:Cu^2+^, (1:1): P<0.05; (1:2): P<0.001; **Fig. 2c, 4b** and **Supplementary Fig. 4c**). Aggregates occurring in [PrP:Cu^2+^+H_2_O_2_] incubations had solvent-exposed hydrophobic residues, revealed by SYPRO orange binding (**Fig. 4c**), and showed intrinsic fluorescence upon excitation in the UV range suggesting amyloid-like cross-β sheet structure^49^ (**Fig. 4d**). After 24 h, amorphous and fibrillar aggregates could be observed (**Supplementary Fig. 4a**). FRAP in HEPES buffer (without salt) showed ~60% recovery for PrP:Cu^2+^ condensates (t1/2 ~13.3 s) versus no recovery for [PrP:Cu^2+^+H_2_O_2_] condensates/aggregates (**Fig. 4e** and **Extended Data Fig. 4b, d**), indicating the loss of molecular diffusion. Notably, PrP:Cu^2+^ recovery was slower than in PhysB (t1/2 ~5.6 s; PhysB contains 120 mM KCl and higher ionic strength, *see* **Supplementary Table 1**) (**Fig. 3a**), suggesting less molecular diffusion inside PrP:Cu^2+^ condensates in the presence of electrostatic interactions (**Extended Data Fig. 4c**).

**Fig. 4.**
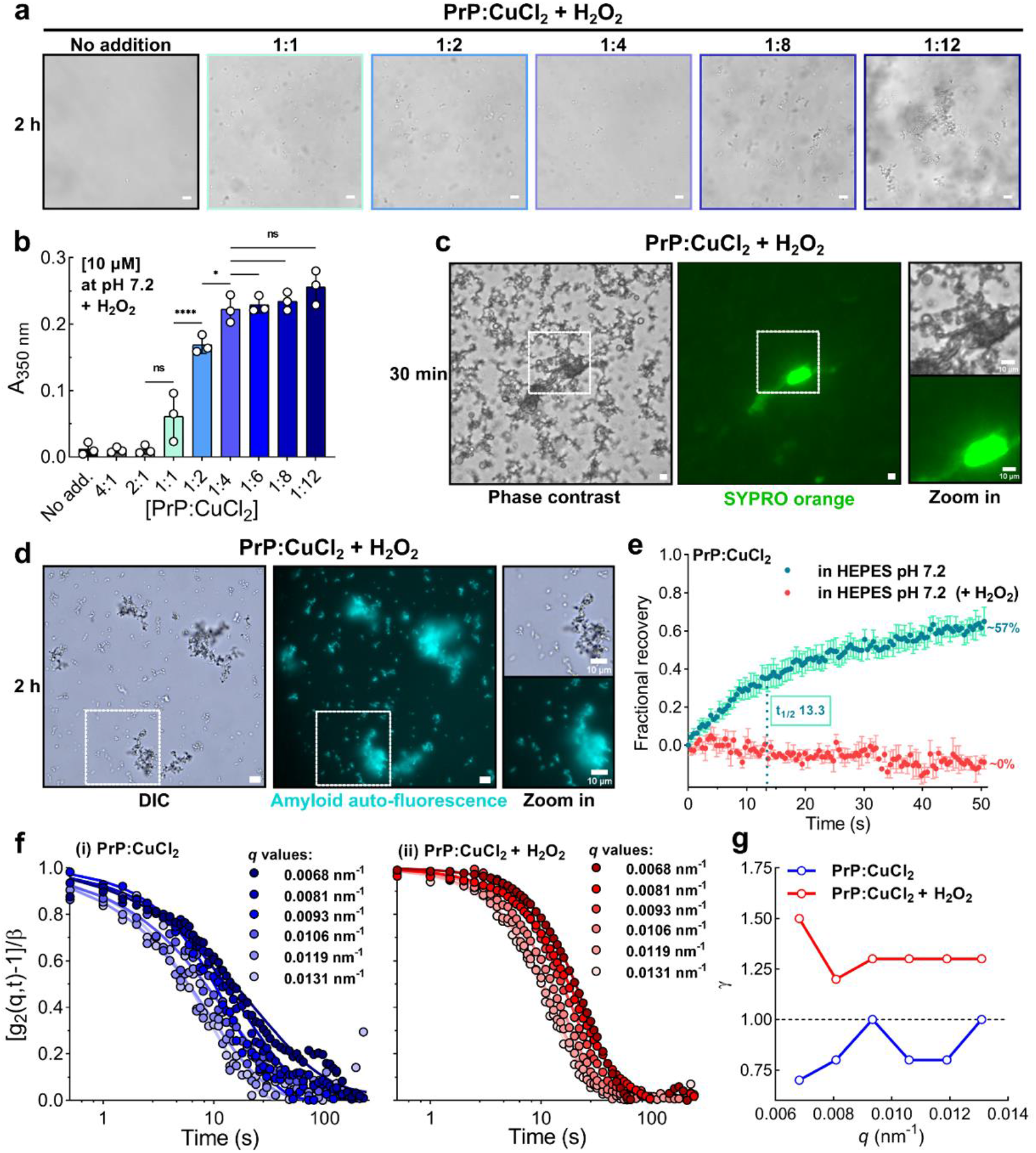
H_2_O_2_ treatment triggers PrP:Cu^2+^ liquid-to-solid phase transition. **a**, Phase contrast images 25 μM PrP incubated with increasing concentrations of CuCl_2_ in the presence of 10 mM H_2_O_2_ (2 h incubation). **b**, Turbidity at 350 nm of 10 μM PrP at increasing CuCl_2_ concentrations (6 minutes after sample preparation) in the presence of H_2_O_2_. Data shown as mean±S.D, n= 3. **c**, Phase contrast (left) and fluorescence (right) images of 25 μM PrP with 200 μM CuCl_2_ and 10 mM H_2_O_2_, stained with SYPRO orange after 30 min (middle). Zooms of ROIs shown on the right. **d**, DIC (left) and fluorescence (right; intrinsic blue fluorescence characteristic of amyloids) images of 25 μM PrP incubated with 200 μM CuCl_2_ and 10 mM H_2_O_2_ after 2 h incubation. Zooms of ROIs shown on the right. **e**, FRAP of 10 μM PrP with 80 μM CuCl_2_ in 20 mM HEPES pH 7.4 with (green; t1/2 is shown in the rectangle) and with the addition of 10 mM H_2_O_2_ (red). FRAP recorded immediately after sample preparation. Data shown as mean ± SEM, n=3 condensates. Scale bars, 10 μm. **f**, XPCS autocorrelation functions for increasing *q* values of PrP:CuCl_2_ (i) and PrP:CuCl_2_+H_2_O_2_ (ii). **g**, XPCS derived Kohlrausch–Williams-Watts exponent, γ, over the recorded *q* range for PrP:CuCl_2_ (blue) and PrP:CuCl_2_+H_2_O_2_ (red). All experiments were performed at 25°C in the PhysB, unless otherwise stated.

### X-ray photon correlation spectroscopy reveals H_2_O_2_-induced differences in PrP:Cu^2+^ condensates

Biomolecular condensates are complex fluids that can become less dynamic over time^50,51^. To gain further insights into the effect of H_2_O_2_ on PrP:Cu^2+^ condensate fluidity, we assessed the dynamic properties of a fluid or polymer on the time (microseconds to hours) and length (nanometers to micrometers) scale relevant for LLPS using X-ray photon correlation spectroscopy (XPCS)^52,53^. Both PrP:Cu^2+^ and [PrP:Cu^2+^+H_2_O_2_] displayed a single exponential decay and dynamics in the range of seconds (**Fig. 4f**). The broad exponential decay of the autocorrelation functions indicated dynamical heterogeneities observed in viscous systems^54^. PrP:Cu^2+^ condensate relaxation dynamics were characterized by sub diffusive motion (γ < 1; for comparison: γ = 1, Brownian motion; γ > 1, super diffusive) over the entire dynamic size scale (acquisition *q*-range; **Fig. 4g** and **Supplementary Fig. 4d**). Slow, sub diffusive relaxation is observed in coarsening viscous condensates^54^, e.g., during LLPS of lysozyme^53^. PrP:Cu^2+^ condensates have characteristics of a viscous fluid that can transition into soft glassy materials, reminiscent of condensates formed by fused in sarcoma (FUS) protein that display an increase in viscosity but no change in elasticity over time, described as aging Maxwell fluids^51^. Similar to PrP:Cu^2+^, FUS condensates do not show changes in morphology during aging, albeit the observed rheological changes (**Supplementary Fig. 4e**). However, when adding H_2_O_2_, we observed remarkable changes in the dynamic properties of condensates (**Fig. 4g**): [PrP:Cu^2+^+H_2_O_2_] condensates/aggregates were characterized by super diffusive (γ > 1) viscoelastic relaxation indicating network fluctuations in the condensates, likely via liquid-to-solid transition during gelation/hardening^52,53^. The loss of molecular diffusion (FRAP data, **Fig. 4e**) and progressive formation of aggregated structures (microscopy data, **Fig. 4a,c,d** and **Supplementary Fig. 4a**) support this idea.

### Cu^2+^ prevents PrP misfolding but in the presence of H_2_O_2_ triggers amyloid formation

To further assess the influence of Cu^2+^ and H_2_O_2_ on PrP aggregation, we performed *in vitro* aggregation assays by seeding PrP with pre-formed recombinant PrP fibrils produced under denaturant condition^55^, in the absence or presence of CuCl_2_ and/or H_2_O_2_ (**Fig. 5a**). This condition would mimic a pathological condition with seeded polymerization of PrP^C^ and oxidative stress (CuCl_2_+H_2_O_2_).

**Fig. 5.**
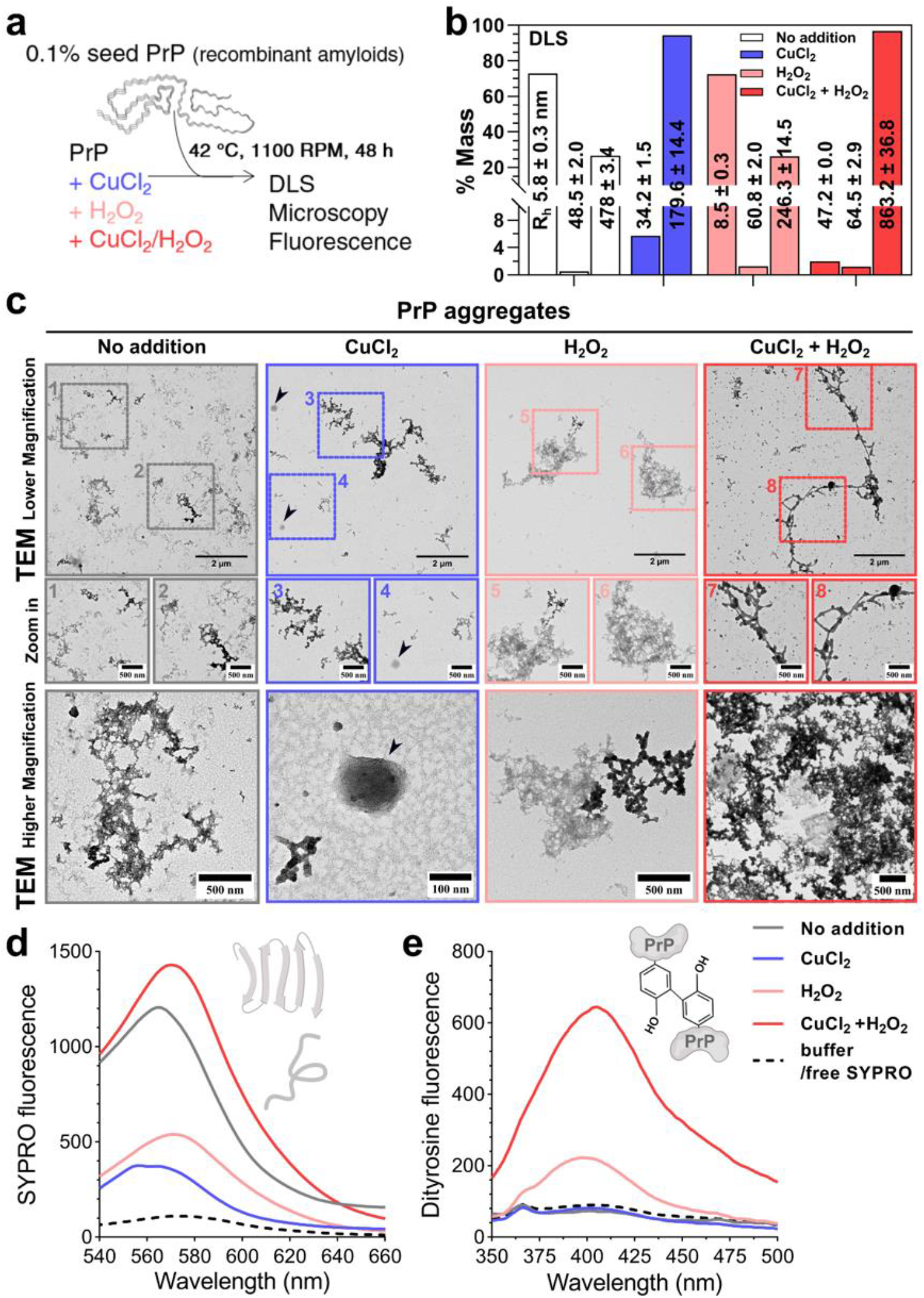
CuCl_2_ reduces the formation of β-sheet-rich structures in PrP aggregates. **a**, Graphic of experimental setup to examine PrP seeded aggregation *in vitro*. PrP (25 μM) was incubated with 0.1% seeds (pre-aggregated PrP obtained from denaturation protocol with 1 M guanidinium-HCl, 3 M urea) under continuous agitation at 42°C for 48 h in the presence or absence of CuCl_2_ (200 μM), H_2_O_2_ (10 mM), or both. **b**, DLS analysis of aggregated 48 h-samples (diluted to 1 μM for measurements). Mass percentage of each species is plotted and R_h_ shown within bar. Data shown as mean ± S.D. from 10 acquisitions at 25°C. **c**, Negative-stain TEM at different magnifications. Arrowheads mark circular condensate-like structures. Zoom-ins from ROIs are marked by numbers. **d**, SYPRO orange emission spectra (excitation 495 nm; emission 535-665 nm) of 48 h-aggregated samples. The illustrations show a β-sheet-rich structure and an unfolded protein which are probed by SYPRO orange. **e**, Dityrosine intrinsic fluorescence (excitation 325 nm; emission 350-500 nm) of 48 h-aggregated samples. Illustration shows an intermolecular DiTyr cross-link. Legend applies to ‘**d**’ and ‘**e**’. All samples were prepared in PhysB.

DLS showed a multimodal particle size distribution for all samples after aggregation, and at least 30% of particles were composed by high-order oligomers (Rh ranging from 180 to 863 nm) (**Fig. 5b**). Of note, monomeric PrP not subjected to the aggregation protocol has a R_h_ of ~2.2 nm (**Fig. 3c**). Addition of CuCl_2_ or CuCl_2_+H_2_O_2_, resulted in the lack of monomeric or low order PrP species (for monomer up to tetramer; R_h_ ranging from 2.2 to 20 nm, considering PrP partially unfolded nature), suggesting a shift to mesoscale and high-order oligomers of PrP. The PrP:H_2_O_2_ sample did not show large particle formation. Oppositely, the PrP:CuCl_2_+H_2_O_2_ sample showed the highest mass of high-order population (Rh >800 nm) (**Fig. 5b**). Therefore, the enhanced phase transition of [PrP:CuCl_2_+H_2_O_2_] may synergize with seeded aggregation of PrP. We then proceeded the characterization of PrP aggregates with transmission electron microscopy (TEM) and thioflavin-T (ThT) fluorescence microscopy. PrP and PrP:H_2_O_2_ formed similar clusters of amorphous aggregates that were positive for ThT (**Fig. 5c**). PrP:CuCl_2_ formed circular structures reminiscent of condensates, few small amorphous aggregates, and diffuse material that may correspond to soluble oligomers. Furthermore, PrP:CuCl_2_ was the only sample lacking thioflavin-T (ThT) positive aggregates (**Supplementary Fig. 5a**). In PrP:CuCl_2_+H_2_O_2_ preparations, we detected large amorphous, branched aggregates as well as long fibrils and clusters of amorphous aggregates with weaved-in circular condensate-like structures.

To explore whether the aggregates observed by TEM contained amyloid-like β-structure, we measured SYPRO orange fluorescence emission, which is ~3-fold more sensitive than ThT for amyloids quantification^56^ (**Fig. 5d**). PrP:CuCl_2_+H_2_O_2_ samples showed the highest SYPRO fluorescence intensity, followed by PrP alone, indicating β-amyloid content in these samples. PrP:CuCl_2_ and PrP:H_2_O_2_ had similarly low SYPRO intensities. These data suggest that H_2_O_2_ promotes PrP amyloid formation in the presence of CuCl_2_, whereas CuCl_2_ alone prevents. Of note, SYPRO fluorescence of PrP:CuCl_2_ showed a 8 nm blue shift of the maximum emission wavelength (λmax=556 nm for PrP:CuCl_2_ vs. 564 nm for PrP), indicating a more hydrophobic environment. Measuring the fluorescence of 1-anilino-8-naphthalenesulfonate (ANS), whose fluorescence intensity is enhanced in hydrophobic environments, confirmed this observation (**Supplementary Fig. 5b,c**). After 48 h of incubation with pre-formed PrP seeds, PrP but not PrP:CuCl_2_ showed enhanced ANS fluorescence (**Supplementary Fig. 5c**), indicating that Cu^2+^ prevented the exposure of hydrophobic residues due to partial unfolding of PrP.

We wondered what would be the mechanism by which CuCl_2_ together with H_2_O_2_ would favor β-aggregation of PrP. Firstly, to determine changes in protein folding that could favor PrP aggregation, we measured intrinsic tryptophan fluorescence during thermal denaturation, right after sample preparation (**Extended Data Fig. 5**). PrP conformation was significantly impacted by CuCl_2_+H_2_O_2_, even at room temperature, evident from a 4-fold decreased fluorescence intensity (vs. PrP; **Extended Data Fig. 5a-d,e**). Additionally, the emission spectrum exhibited a red shift (**Extended Data Fig. 5d,f**), possibly due to dityrosine formation (emission at 405 nm). PrP β-aggregation may, thus, be promoted by unfolding and the generation of dityrosine crosslinks due to oxidation in the presence of CuCl_2_+H_2_O_2_.

In metal-catalyzed oxidation conditions (e.g., CuCl_2_+H_2_O_2_), covalent intra- and intermolecular cross-links that promote protein aggregation can occur through dityrosine (DiTyr) formation^57^. We investigated whether H_2_O_2_ would trigger DiTyr formation in PrP, which is apparent from an increase in DiTyr fluorescence. Indeed, the presence of CuCl_2_+H_2_O_2_ significantly increased DiTyr formation in PrP aggregates compared to all other conditions, *e.g.*, 3-fold increased emission at 405 nm compared to PrP:H_2_O_2_; **Fig. 5e**). Eleven out of 13 tyrosine residues are in the C-terminal half of PrP that forms the protease and denaturation resistant-core of pathological PrP aggregates^1^, likely explaining why oxidation-induced DiTyr cross-linking favors PrP aggregation.

In addition, ROS produced by Fenton reactions as catalyzed by Cu^2+^+H_2_O_2_, induce pathological ‘β-cleavage’ of PrP^C^ around residue 90, releasing the N-terminal part extracellularly while keeping the C-terminal part attached to the cell surface^58,59^. We evaluated whether recombinant PrP with CuCl_2_+H_2_O_2_ would undergo β-cleavage *in vitro* (**Supplementary Fig. 6**). SDS-PAGE showed that PrP:CuCl_2_+H_2_O_2_ samples contained full-length PrP (~22 kDa) and SDS-stable higher-order oligomers right after sample preparation (**Supplementary Fig. 6b**, first lane). After 30 min, these PrP species started to disappear and a 15-20 kDa fragment appeared (~16.3 kDa; arrow; **Supplementary Fig. 6b**), likely corresponding to the C-terminal product of β-cleavage. After 120 min, ~75% of full-length PrP and the C-terminal fragments were degraded in [PrP:CuCl_2_+H_2_O_2_] (**Supplementary Fig. 6c**).

In summary, the presence of copper in oxidative conditions (CuCl_2_+H_2_O_2_) seems to trigger PrP aggregation via multiple mechanisms, including PrP unfolding and exposure of hydrophobic residues, production of aggregation prone C-terminal fragment through β-cleavage, and the formation of DiTyr crosslinks.

### Extended exposure to Cu^2+^ induces intracellular PrP aggregation

We asked whether CuCl_2_ and ROS would also induce PrP aggregation in cells, for example, promoted by PrP oxidation and/or β-cleavage. Because cellular systems naturally possess H_2_O_2_ and other ROS as metabolism by-products, Cu^2+/+^ can catalyze further ROS production by intra- and extracellular Fenton reactions. We incubated HEK293 cells expressing PrP^C^-YFP-GPI with 300 μM CuCl_2_ for 3 h (of note, experiments presented in **Fig. 1** were conducted within 1 h) and tested the formation of amyloid-like aggregates by addition of amyloid dyes. Indeed, Cu^2+^ exposure for 3 h led to AmyTracker680-positive aggregates at the cell surface (**Fig. 6a,b**). In addition, we observed the formation of vesicle-like hollow, spherical structures in the cytosol that carried PrP^C^-YFP-GPI in their surrounding membrane (**Fig. 6c** and **Supplementary Video 4**). We speculate that these might be stress-induced ER vesicles^60^. When incubating cells with CuCl_2_ in serum-free imaging solution (lacking ROS scavenging proteins like albumin^61^), we observed PrP^C^-YFP-GPI signal near the nucleus and Nile Red-positive PrP aggregates **(Supplementary Fig. 7a,b**). Notably, HeLa cells transfected with PrP^C^-YFP-GPI incubated with CuCl_2_ showed hollow spherical ER-like structures already after 1 h, indicating that these cells may be more vulnerable to PrP:CuCl_2_ induced conditions (**Supplementary Fig. 7c)**. These data corroborate the observed amyloid-like aggregates of [PrP:CuCl_2_+H_2_O_2_] (**Fig. 4** and **Fig. 5**) indicating that oxidative stress promotes PrP liquid-to-solid transition *in vitro* and *in celullo*.

**Fig. 6.**
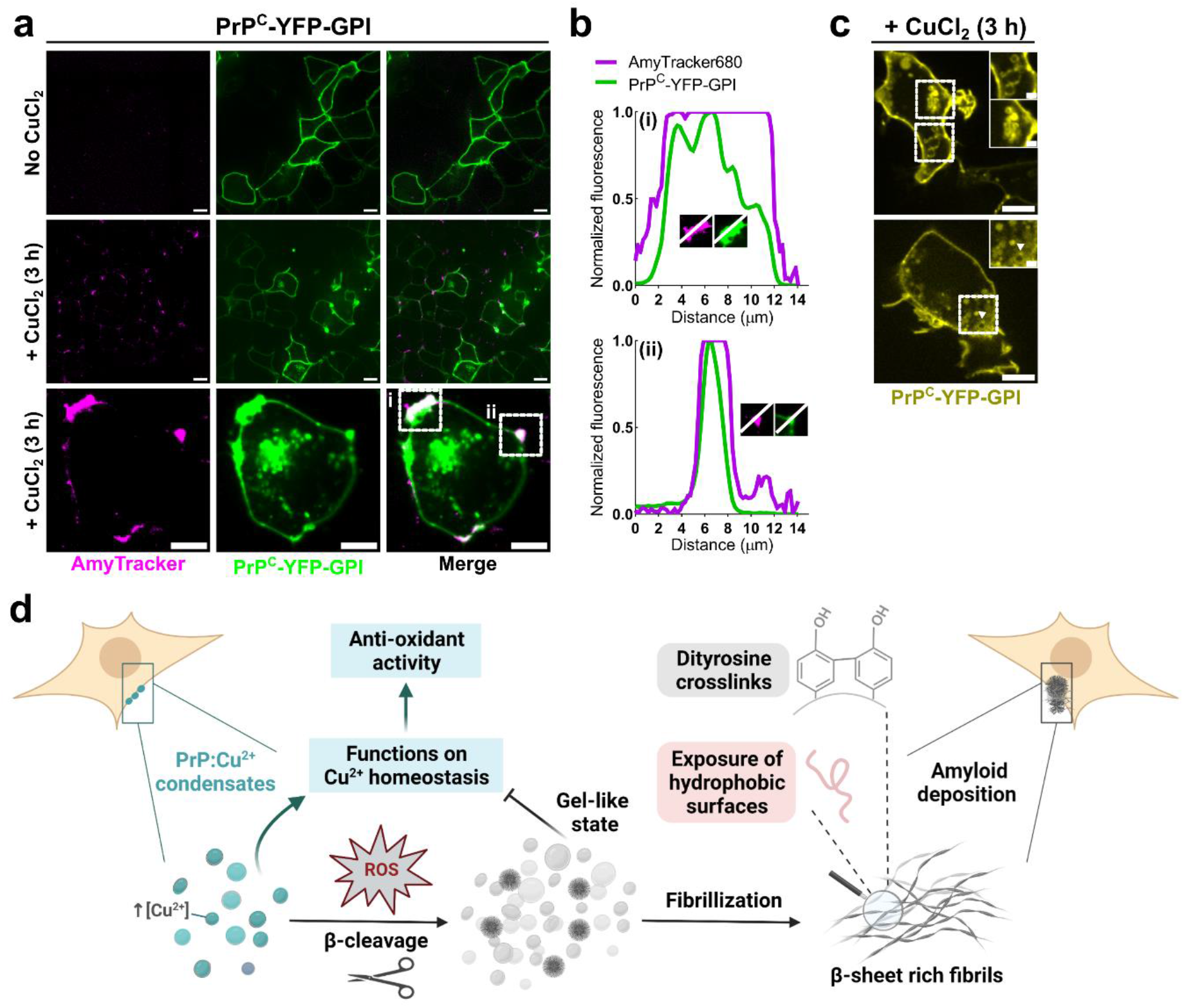
Long-term exposure to Cu^2+^ leads to PrP^C^-YFP-GPI amyloid-like aggregates in cells. **a**, Live cell imaging of HEK293 cells transfected with PrP^C^-YFP-GPI. Cells treated with 300 μM CuCl_2_ for 3 h (middle and bottom row) showed AmyTracker680 positive aggregates on the cell surface. Bottom: individual cell with PrP agrgegates. White lines indicate positions of line profiles shown in (**b**). **b**, line profiles through AmyTracker positive PrP aggregates on the surface of the cell shown in (**a**, bottom). **c**, Formation of spherical hollow structures with PrP^C^-YFP-GPI in the surrounding membrane after 3 h Cu^2+^-treatment. **d**, Model for the role of copper in PrP^C^ condensation and oxidation-induced aggregation. Left: PrP:Cu^2+^ condensation on the cell surface buffers Cu^2+^ concentration by sequestering redox-active excessive copper. This reduces the oxidative burden and prevents cellular damage. In the presence of ROS, further stimulated through Fenton reactions of Cu^2+^, PrP:Cu^2+^ condensates transition into a gel-like state (less molecular mobility inside condensates) that inhibits functional Cu^2+^ sequestering by PrP. Moreover, ROS triggers the unfolding of PrP resulting in exposure of hydrophobic surfaces, drives PrP β-cleavage that produces aggregation-prone C-terminal PrP fragments, and enhances DiTyr crosslinking, altogether driving PrP aggregation into amyloid-like aggregates. In summary, PrP controls copper homeostasis and prevents ROS generation through LLPS, but abnormally high or prolonged ROS can result in aberrant PrP condensation and pathological aggregation.

## Discussion

In the last 25 years, it became clear that PrP plays a key role on copper homeostasis^10,12^, and that, in turn, copper influences PrP aggregation^14,62,63^. With our data, we now provide mechanistic evidence that both of these observations are related to PrP LLPS and its modulation by Cu^2+^. Earlier works showed PrP (protein domains or mutants) LLPS *in vitro* in simple buffer systems and without consideration of physiological Cu^2+^ concentration^25,26,42,64–66^. By investigating the effects of Cu^2+^ on full-length PrP condensation and aggregation at physiologically relevant PrP concentrations (4 μM^25^), ionic strength, pH, and molecular crowding, we not only complement previous studies, but are able to provide evidence for the role of PrP:Cu^2+^ condensation in the cellular context.

From our data, we conclude that Cu^2+^ binding to His residues in the octapeptide region of PrP, the main copper-binding site, induces structural changes that promote multivalent molecular interactions of PrP molecules driving its condensation. Such interactions could, for example, induce LARKS motifs reported to promote protein LLPS^20^, supporting previous reports on PrP N-terminal domain enhanced structural stability in the presence of Cu^2+^ ^13,63^. Previously reported copper-mediated interactions between N- and C-terminal PrP domains aiding protein compactness^13,17^ could be involved as well. Importantly, reducing Cu^2+^ binding by EDTA or low pH reversed PrP LLPS, and Cu^2+^ prohibited PrP aggregation, suggesting that PrP:Cu^2+^ LLPS and aggregation may occur independently, similarly to Tau^67^, or even that PrP:Cu^2+^ condensates might suppress aggregation as shown for α-synuclein^68^. Interestingly, the octarepeats region in PrP is conserved across mammalian and avian species^69^, indicating that Cu^2+^-driven PrP LLPS may play a crucial role for cellular copper homeostasis.

In cells overexpressing PrP^C^, Cu^2+^ induced large plasma membrane-bound PrP^C^ clusters that showed characteristics of liquid-like condensates (*i.e.*, reduced molecular diffusion, fission, fusion). Absorption of Cu^2+^ in cell surface PrP^C^ condensates at a 4:1 (Cu^2+^:PrP) stoichiometry, similar to our *in vitro* observations, would provide an effective Cu^2+^ buffering system that counteracts metal ion toxicity, before redox-active copper would enter the cytosol. Furthermore, lysosomal storage of Cu^2+^, and its redistribution to cellular compartments and metalloproteins that ‘quench’ Cu^2+^ redox-activity^70^, could be facilitated through internalization of plasma membrane PrP^C^:Cu^2+^ condensates^71^ by the endolysosomal pathway. In this case, we could expect release of Cu^2+^ in the acidic environment inside lysosomes (*in vitro:* no LLPS and no Cu^2+^ binding at pH~5), which would enable lysosomal Cu^2+^ storage and the recycling of PrP^C^ back to the plasma membrane, to sequester more Cu^2+^ through co-condensation. In fact, previous works suggested that PrP^C^ may act as copper buffer protecting cells from oxidative burden through an unknown mechanism, for example, during neuronal depolarization^12^. Our data delivers a LLPS-based cellular mechanism for these observations, in which PrP^C^ condensates sequester copper ions and prevent oxidative burden. Notably, PrP LLPS was also triggered by other metal ions *in vitro* (**Extended data Fig. 3**), suggesting that membrane-standing PrP^C^ condensates could provide a similar mechanism for other metal ions as well. Dysregulation of PrP:metal ion co-condensation could result in metal dyshomeostasis, as it is observed early onset prion disease^14^. Interestingly, PrP^C^:Cu^2+^ condensates formed preferentially at cell-cell junctions, where PrP:Cu^2^’ condensation could also be involved in copper exchange between cells. PrP was reported to interact with proteins in adherens junctions and desmosomal junctions^72,73^, and we recently suggested that the multivalency of these interactions could result in LLPS at cell-cell contacts^29^.

Long Cu^2+^ incubation led to amyloid-like PrP^C^ aggregation on the cell surface, which is in line with the proposed pathological PrP^C^ to PrP^Sc^ conversion at the plasma membrane in prion diseases^2^. From our *in vitro* data, we would assign this observation to the presence of ROS in the culture medium. ROS formation, triggered by Cu^2+^-catalyzed Fenton reactions, resulted in *i*, PrP β-cleavage, releasing the LLPS driving N-terminal domain of PrP^74^ and producing the aggregation prone C-terminal fragment in minutes; *ii* triggered DiTyr crosslinking, and *iii*, induced liquid-solid transition of PrP condensates and the formation of amyloid-like PrP aggregates. In contrast, in non-oxidizing conditions (no H_2_O_2_), Cu^2+^ prevented PrP aggregation, supporting earlier findings that PrP:Cu^2+^ aggregates do not have amyloid content^62,75^.

Taken together, our data provide mechanistic insights into Cu^2+-^driven PrP condensation as a way of cellular copper ion buffering and reveals the effects by which ROS triggers PrP aggregation in the presence of copper (**Fig. 6d**).

## Abbreviations

CD: circular dichroism
DIC: differential interference contrast
DLS: dynamic light scattering
FRAP: fluorescence recovery after photobleaching
LLPS: liquid-liquid phase separation
PM: plasma membrane
PrP: prion protein
XPCS: x-ray photon correlation spectroscopy

## Methods

### Recombinant protein production

pET-41 containing the coding region for mature full-length mouse PrP (residues 23-231) was transformed in *Escherichia coli* BL21. Protein expression was induced at 37 °C for 16-18 h in Luria-Bertani medium (LB) by addition of 0.5 mM isopropyl thio-β-D-galactoside (IPTG), followed by PrPpurification as previously described^76^. Briefly, purification was achieved using Ni^2+^-affinity denaturing chromatography followed by in-column refolding and dialysis against ultrapure water. All chromatography steps were performed in an AKTA prime liquid chromatography system (GE Healthcare). Aliquots of PrP were kept at −20 °C for up to 6 months.

### AlexaFluor-647 labelling of PrP

Purified PrP (150 μM; 0.5 mL) in 0.1 mM sodium bicarbonate pH 8, 1 M NaCl was incubated with 100 μg succinimidyl ester of AlexaFluor-647 (ThermoFisher, A20006) in the dark for 1 h at 25°C with gentle shaking. Uncoupled fluorophores were removed by chromatography in a HiTrap desalting column (Cytiva) equilibrated in 10 mM sodium phosphate pH 5.8. The degree of labeling was calculated using dye absorbance at 650 nm and a molar extinction coefficient of 239,000 cm^-1^ M^-1^. We obtained a PrP (2 μM; 3 mL) solution with a labeling yield of 1.2 moles of AlexaFluor-647 per mole of PrP. The low protein yield is due to PrP precipitation and aggregation at basic pH.

### Sample preparation for LLPS assays

Samples of 25 μM PrP were immediately suspended in 0.22 μm-filter sterilized physiological-like buffer (**Table 1**). Subsequently, samples were incubated with 25 to 600 μM CuCl_2_, and/or 10 mM H_2_O_2_, a known oxidizing conditions for PrP, as reported in ^77,78^. CaCl_2_ was the first to be dissolved whereas K_2_HPO_4_/KH_2_PO_4_ was the last one added, to prevent Ca_3_(PO_4_)_2_ precipitation, followed by addition of PEG4K and pH adjustment with KOH. The buffers were prepared 2x concentrated and kept at 4 °C for one month. In case of turbidity measurements, the concentration of PrP samples were 10 μM in the presence or not of 2.5 to 120 μM CuCl_2_ and/or 10 mM H_2_O_2_.

### *In vitro* liquid-liquid phase separation studies by microscopy and turbidity

96-well plates (flat bottom, Sarstedt Inc #82.1581) were coated with 0.5% (w/v) bovine serum albumin (BSA, Sigma-Aldrich A2153) in 1x phosphate-buffered saline (PBS) for 1 hour in a see-saw rocker followed by 2× rinsing with ultrapure water. This step was carried out to provide a hydrophilic surface to the polystyrene plates. Protein solutions were immediately prepared in specified buffers as indicated in figure legends and 50 μL (imaging) or 80 μL (turbidity) solutions were added to each well followed by a 30 min incubation at R.T. Imaging was carried out immediately after reactants addition (specified transition metals or CuCl_2_ and/or H_2_O_2_) or at specified times in the figures. Samples described above were imaged in an inverted phase contrast microscope (EVOS® FL Cell Imaging System, ThermoFisher) with a 40 × apochromat objective. For Zincon staining, phase separated samples were deposited onto poli-D-lysine coated glasses and incubated for 30 minutes. Followed by the addition of a 0.4 mM Zincon solution (stock dissolved in 15 mM Tris-HCl pH 7.4) and mount on coverslips sealed by nail polish. Turbidity (detected as absorbance at 350 nm) assays were performed in a Synergy H1 spectrophotometer (BioTek Instruments) at 28 °C.

### Spectrophotometric determination of Cu^2+^ in the dense phase

A solution of PrP at 25 μM in 50 mM HEPES pH 7.4, 125 mM NaCl, 150 μM CaCl_2_, 5 mM MgCl_2_, 6% (w/v) PEG4K was incubated with 8 x molar excess of CuCl_2_ (200 μM). Upon addition of CuCl_2_, phase separation was triggered, followed by sample equilibration for 1 h in an ice bath at 0°C. To spin down the droplets, the sample was centrifuged at 100 x g for 30 minutes in a temperature-equilibrated centrifuge at 4 °C. A blue pellet (*i.e*. dense phase) was formed in the bottom, all supernatant (*i.e*. light phase) was removed to another tube. The pellet was weighted (0.0010 g) and solubilized in 4 μL of the light phase with addition of urea and nitric acid to a final concentration of 4 M and 2.4%, respectively, totalizing 20 μL of the pellet-containing solution. For colorimetric quantification of Cu^2+^ we used a previously published protocol^38^ in which samples are prepared in borate buffer containing 8 M urea (to release Cu^2+^ from protein).

### Fluorescence recovery after photobleaching

*In vitro* FRAP experiments were performed in a confocal microscope (LSM 710, Zeiss) using a 100 × oil immersion objective and 3× digital zoom. Solutions of unlabeled PrP spiked with 0.1% PrP-Alexa647 were imaged (excitation wavelength at 633 nm; emission 638-755 nm) in a confocal dish (SPL Life Sciences, 200350). Bleaching was performed with a 405 nm laser (100% intensity, 1000 iterations; ~14 s total time) was started after 10 scans (scan rate 390 ms) in circular ROIs (0.8-1.0 μm diameter) and monitored for 50-90 seconds. FRAP experiments in HEK293 cells (transfected with PrP^C^-YFP-GPI) were performed in a spinning disk confocal microscope (Eclipse-Ti CSU-X, Nikon) equipped with an incubator for live cell imaging (37 °C; 5% CO_2_) using a 60 × oil immersion objective. After 5 frames (scan rate 200 ms), bleaching was set to 2 loops with 10% 488 nm laser and the recovery was monitored for 120 s. In all cases, unbleached and backgrounds ROIs of same size (4.6 μm^2^) were acquired simultaneously. Data were background subtracted and normalized to the reference ROI to account for photofading during acquisition.

### AmyTracker and Nile Red microscopy

*In vitro* LLPS with AmyTracker was conducted with 1:5000 AmyTracker 680. Labelling of living cells was performed with 1:1000 AmyTracker 680 in Opti-MEM for 30 min followed by replacing with fresh medium containing CuCl_2_ or not. In case of Nile Red staining (0.1 μg/mL), incubation was carried out in imaging solution (20 mM HEPES pH 7.4, 140 mM NaCl, 2.5 mM KCl, 1.8 mM CaCl_2_, 1.0 mM MgCl_2_, 20 mM D-glucose) for 30 min followed by replacing with fresh medium containing CuCl_2_ or not.

### Transmission electron microscopy

Micrographs of PrP in the absence or presence of CuCl_2_, H_2_O_2_ or both were acquired on a Hitachi HT7800 electron microscope operating at 100 kV. Grids were prepared by spotting 5 μL of samples on formvar/carbon-coated grid followed by contrasting with 2.0% uranyl acetate solution.

### Turbidity as a function of temperature

Absorbance readings at 350 nm were carried out in a V-730 UV-VIS spectrophotometer (Jasco Corporation) equipped with a Koolance Cooling System EX2-755 using the Temperature Interval Scan Measurement tool. 60 μL of samples in 3 mm quartz cuvettes with corresponding micro cell jackets. Samples were previously equilibrated at the starting temperature for 30 minutes before measurements. A temperature gradient from 38 to 14 °C and the return to the start temperature was run with a rate of 0.5 °C/min and incubated for 30 seconds before each read.

### Dynamic Light Scattering (DLS)

Reads (10 accumulations each) were collected in a DynaPro NanoStar equipment (Wyatt Technology, CA, USA) with a GaAs laser at λ=658 nm and intensity from 100 mW. Samples were immediately prepared in the physiological-like buffer pH 7.2. The hydrodynamic radius of PEG4K in buffer solutions (without protein) was monitored and showed monodispersity, with particles varying from 0.7-0.9 nm and in all reads corresponded to 100% mass. DLS data were corrected with buffer properties, *i.e*. physiological-like buffer refraction index and viscosity at specified temperatures. The physiological-like buffer refraction index at 25°C (1.342) was measured using an Abbe refractometer. This buffer contains 6% (w/v) PEG4K, thus, to determine its viscosity (η), we retrieved the viscosity data of PEG4K aqueous solutions at 25°C determined by ^79^. From the linear regression, we obtained the viscosity of the physiological-like buffer (1.982 × 10^3^ PL).

### Cell culture, transfection, and cell viability analysis

HEK293 cells were cultured in RPMI medium (Sigma) supplemented with 10% v/v fetal bovine serum (FBS) and 1% antibiotics penicillin/streptomycin (Gibco) at 37 °C in 5% v/v CO_2_ atmosphere. For cell transfection, cells were plated in 24-well plates (viability assay), 8-well confocal dishes (imaging and FRAP) and transfected after 24 hours with Lipofectamine 2000 (ThermoFisher) using 400 ng-1 μg of specified plasmids (YFP-GPI^80^, PrP^C^-YFP-GPI or GFP) in Opti-MEM (ThermoFisher). Following 48 hours post transfection, cells were treated with 10-1000 μM of dihydrated CuCl_2_ (Sigma) for 1 hour. After treatment, cells were detached with trypsin solution (Nova Biotecnologia) and quantified using Trypan blue solution (0.2% w/v in PBS) in a hemocytometer. For FRAP, transfected cells had medium exchanged to fresh Opti-MEM 1 h before addition of 300 μM CuCl_2_. After 30 min to 1 h incubation with CuCl_2_ or not, FRAP experiments were performed.

### Immunoblotting

Whole cell extracts were obtained 24 hours post transfection using mild RIPA buffer (50 mM Tris pH 7.4, 150 mM NaCl,1 mM EDTA and NP40 1% v/v) supplemented with protease inhibitor cocktail l% v/v (Sigma) and quantified by Bradford assay. Protein extracts (50 μg/lane) were resolved in 15% SDS-PAGE followed by immunoblotting using anti-GFP (AB3080P, Merck), anti-PrP^C^ monoclonal antibody (anti-CD320 clone 6D11; Biolegend, 808001), and anti-β-actin (Invitrogen, MA5-15739) as loading control. Horseradish peroxidase (HRP)-linked conjugated secondary antibodies (Invitrogen) were developed with SuperSignal West Pico Chemiluminescent Substrate (ThermoScientific, 34077) using a Chemidoc scanner (Bio-Rad).

### X-ray photon correlation spectroscopy

XPCS experiments were performed at the CATERETÊ beamline (Brazilian Center for Research in Energy and Materials (CNPEM)) at a photon energy of 10 keV (l% bandwidth). Data were collected using a PiMega540D detector in small angle x-ray scattering (SAXS) geometry at a sample-detector distance of 27.6 m allowing measurement of low scattering vector (*q*) values (*q* = 4πsinθ/λ, being 2θ the angle between the incident and scattered beam). Samples of PrP (25 μM) with or without CuCl_2_ (200 μM) and/or H_2_O_2_ (10 mM) were prepared immediately before measurements and condensates formation was monitored by DIC microscopy *in situ* (Leica LMD 7). Each sample was filled into 1.5 mm diameter quartz capillaries. To mitigate radiation damage, samples were measured with sapphire attenuator and the total time was considered as time with no visible change in the SAXS patterns. The XPCS experiments have been performed below this threshold dose value. All samples were analyzed in the physiological-like buffer pH 7.2 at 25°C. The autocorrelation functions can be expressed by the Kohlrausch-Williams-Watts (KWW) model (equation 1) containing information about the time scales and type of motion in protein condensates.

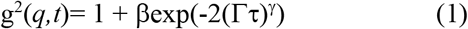

where β is the contrast factor, Γ is the relaxation rate, τ is the delay time, *q* is the scattering vector, and γ is the KWW exponent. The dynamics is quantified by the relaxation rate (**Supplementary Fig. 7d**) and characterized by the KWW exponent γ (**Fig. 4h**), where γ = 1 represents a single exponential decay corresponding to Brownian diffusion. When γ >1 the relaxation process is faster than exponential (super diffusive), often seen in disordered soft solids like gels, and γ < 1 is indicative of relaxation slower than exponential decay (sub diffusive), commonly observed in systems with several competing relaxation mechanisms, such as liquids near the glass transition temperature.

### Steady-state fluorescence and anisotropy

Intrinsic tryptophan fluorescence was recorded with excitation set at 295 nm and emission verified from 310 to 400 nm with 5 μM PrP and varying CuCl_2_ concentration (from 5 to 40 μM). Tryptophan fluorescence as a function of temperature was collected upon slowly heating in steps of 1°C from 25°C to 80°C. The center of spectral mass (CM) in nm was calculated via the equation: CM = Σ νiFi/Σ Fi, where vi is the wavelength and Fi is the fluorescence intensity at each wavelength. Dityrosine (DiTyr) fluorescence was recorded with excitation at 325 nm and emission from 350 to 500 nm. SYPRO orange fluorescence (stock 5000 x in DMSO; working concentration 50 x) was recorded with excitation at 495 nm and emission from 535 to 665 nm. Both DiTyr and SYPRO fluorescence were monitored for 25 μM PrP aggregated samples. The probe 1-anilinonaphthalene-8-sulfonic acid (1,8-ANS at 30 μM) was added to aggregated or soluble 20 μM PrP, and its fluorescence emission was collected from 400 to 600 nm upon excitation at 360 nm. Tryptophan anisotropy (excitation 280 nm; emission 353 nm) of 25 μM PrP was acquired upon CuCl_2_ titration at 25°C in a Cary Eclipse Fluorescence Spectrophotometer (Agilent) with slits of 5 nm (excitation) and 10 nm (emission). All assays were performed in the physiological-like buffer pH 7.2 at 25°C in a FP-8250 Spectrofluorometer (Jasco) equipped with a thermal-control unit (with a Koolance Cooling System EX2-755), unless otherwise stated.

### Circular dichroism

CD data were collected on a Chirascan Circular Dichroism Spectrometer (Applied Photophysics) using 2 mm quartz cell. Spectra were recorded over 190 to 260 nm wavelength range at 1 nm intervals at a speed of 0.5 s per point and reported as the average of 3 measurements. All spectra were subtracted from the corresponding background buffer spectrum and smoothed by a Savitzky-Golay filter with window size of 4. Data were expressed as molar ellipticity [θ] or ellipticity (θ) in millidegrees (mdeg). The mean amino acid residue weight (MRW) of PrP is 100. Molar residue ellipticity [θ] was calculated with the following equation: [θ] = (MRW × θλ)/ (c × l × 10) where θλ is the observed ellipticity (millidegrees), c is the protein concentration (g/mL), and l is the cell pathlength (0.2 cm). Spectra collected in 2 × diluted PhysB (5 mM K_2_HPO_4_/KH_2_PO_4_ pH 7.2, 60 mM KCl, 2.5 mM NaCl, 75 μM CaCl_2_, 2.5 mM MgCl_2_ 3% (w/v) PEG4000).

### Image, data, and statistical analysis

Phase contrast, DIC and fluorescence representative micrographs from at least three independent experiments had their brightness/contrast corrected by histogram stretching using Fiji^81^. Line profile, image quantification and in cell FRAP were analyzed using Fiji. All data and statistical tests (specified in the figure legends) were plotted and analyzed using Prism 8.1.1 (GraphPad Software).

### Data availability

All data are available in the main text or the Supplementary Material. Expression plasmids and cell lines are available upon request.

## Acknowledgements

We thank CENABIO (Universidade Federal do Rio de Janeiro, Brazil) and AMBIO (Charité Universitätsmedizin, Berlin) imaging facilities. We thank Dr. Marilene H. Lopes (Universidade de São Paulo, Brazil) for providing us with the PrP^c^-YFP-GPI plasmid and Dr. Daniel Legler (Universität Konstanz, Germany) for the YFP-GPI plasmid. We thank Dr. Jerson L. Silva (UFRJ, Brazil) for insightful discussions. We thank Sabrina Hübschmann (DZNE-Berlin, Germany) and Maria Heloisa Freire (UFRJ, Brazil) for their help in cell culture. We thank Thayna Sisnande (UFRJ, Brazil) for help in imaging Zincon stained samples. Figure 6d was created with BioRender. Funding: This study was financed by the Coordenação de Aperfeiçoamento de Pessoal de Nível Superior - Brasil (CAPES) (Finance Code 001) (M.J.A.), Fundação de Amparo à Pesquisa do Estado do Rio de Janeiro (FAPERJ) (Y.C. 202.625/2019, 200.562/2023, R.S.C. 201.439/2021, A.P. 200.977/2022, and M.S.A.), the Conselho Nacional de Desenvolvimento Científico e Tecnológico (CNPq) (Y.C. 307294/2018-8, A.P. 306710/2021-8, and M.S.A.), the Brazilian Synchrotron Light Laboratory (LNLS) of the Brazilian Ministry for Science, Technology, Innovations and Communications (MCTIC) (CATERETE-20220546) (Y.C.). M.J.A. is grateful to the CAPES-PrInt scholarship (grant number 88887.695190/2022–00). S.W. provided funding through German Research Society (DFG) through the priority program SPP2191 (419138680) and the Hertie Foundation (P1200002).

## Author contributions

M.J.A. conceived and designed the study, wrote the original draft, involved in all experiments and data analysis. A.R.P. and Y.C. performed the XPCS data collection and analysis. S.M. performed live-cell imaging and FRAP, helped with image analysis. T.S.L. and R.C.S. performed cell viability assay, western blotting and analysis. Y.C., S.W., M.S.A., A.S.P., R.C.S., grant acquisition. Y.C., M.S.A., A.S.P., S.W., supervised the study. Y.C., A.S.P., M.S.A., edited the first draft. S.W., edited the manuscript to the final version. All authors contributed to the article and approved the manuscript.

## Competing interests

The authors declare no competing interests.

## Additional information

Extended data

Supplementary Material

## Notes

### Competing Interest Statement

The authors have declared no competing interest.

## References

1. Prusiner, S. B. Prions. Proc. Natl. Acad. Sci. 95, 13363–13383 (1998).

2. Caughey, B., Baron, G. S., Chesebro, B. & Jeffrey, M. Getting a grip on prions: oligomers, amyloids, and pathological membrane interactions. Annu. Rev. Biochem. 78, 177–204 (2009).

3. Kim, S. J. & Hegde, R. S. Cotranslational partitioning of nascent prion protein into multiple populations at the translocation channel. Mol. Biol. Cell (2002) doi: 10.1091/mbc.E02-05-0293.

4. Johnson, C. J. et al. Low copper and high manganese levels in prion protein plaques. Viruses (2013) doi:10.3390/v5020654.

5. Milhavet, O. et al. Prion infection impairs the cellular response to oxidative stress. Proc. Natl. Acad. Sci. U. S. A. (2000) doi:10.1073/pnas.250289197.

6. Pushie, M. J. et al. Prion protein expression level alters regional copper, iron and zinc content in the mouse brain. Metallomics (2011) doi:10.1039/c0mt00037j.

7. Nishimura, T. et al. Cellular prion protein regulates intracellular hydrogen peroxide level and prevents copper-induced apoptosis. Biochem. Biophys. Res. Commun. (2004) doi:10.1016/j.bbrc.2004.08.087.

8. Brown, D. R., Schulz-Schaeffer, W. J., Schmidt, B. & Kretzschmar, H. A. Prion protein-deficient cells show altered response to oxidative stress due to decreased SOD-1 activity. Exp. Neurol. (1997) doi:10.1006/exnr.1997.6505.

9. Millhauser, G. L. Copper and the prion protein: methods, structures, function, and disease. Annu.Rev.Phys.Chem. 58, 299–320 (2007).

10. Brown, D. et al. The cellular prion protein binds copper in vivo. Nature 390, 684–687 (1997).

11. Walter, E. D., Chattopadhyay, M. & Millhauser, G. L. The affinity of copper binding to the prion protein octarepeat domain: Evidence for negative cooperativity. Biochemistry (2006) doi:10.1021/bi060948r.

12. Salzano, G., Giachin, G. & Legname, G. Structural Consequences of Copper Binding to the Prion Protein. Cells 8, 770 (2019).

13. Hodak, M., Chisnell, R., Lu, W. & Bernholc, J. Functional implications of multistage copper binding to the prion protein. Proc. Natl. Acad. Sci. U. S. A. (2009) doi:10.1073/pnas.0903807106.

14. Brown, D. R. Metals and Prions: Twenty Years of Mining the Awe. in Biometals in Neurodegenerative Diseases: Mechanisms and Therapeutics (2017). doi:10.1016/B978-0-12-804562-6.00007-5.

15. Brown, D. R., Clive, C. & Haswell, S. J. Antioxidant activity related to copper binding of native prion protein. J. Neurochem. (2001) doi: 10.1046/j.1471-4159.2001.00009.x.

16. Millhauser, G. L. Copper Binding in the Prion Protein. Acc. Chem. Res. (2004) doi:10.1021/ar0301678.

17. Schilling, K. M. et al. Both N-Terminal and C-Terminal Histidine Residues of the Prion Protein Are Essential for Copper Coordination and Neuroprotective Self-Regulation. J. Mol. Biol. 432, 4408–4425 (2020).

18. Watanabe, Y. et al. A novel copper(II) coordination at His186 in full-length murine prion protein. Biochem. Biophys. Res. Commun. (2010) doi:10.1016/j.bbrc.2010.03.003.

19. Martin, E. W. et al. Valence and patterning of aromatic residues determine the phase behavior of prion-like domains. Science 367, 694–699 (2020).

20. Hughes, M. P. et al. Atomic structures of low-complexity protein segments reveal kinked b sheets that assemble networks. Science 359, 698–701 (2018).

21. Boeynaems, S. et al. Protein Phase Separation: A New Phase in Cell Biology. Trends Cell Biol. 28, 420–435 (2018).

22. Alberti, S. & Hyman, A. A. Biomolecular condensates at the nexus of cellular stress, protein aggregation disease and ageing. Nat. Rev. Mol. Cell Biol. 22, 196–213 (2021).

23. Elbaum-Garfinkle, S. Matter over mind: Liquid phase separation and neurodegeneration. The Journal of biological chemistry (2019) doi:10.1074/jbc.REV118.001188.

24. Wegmann, S. et al. Tau protein liquid–liquid phase separation can initiate tau aggregation. EMBO J. (2018) doi:10.15252/embj.201798049.

25. Kostylev, M. A. et al. Liquid and Hydrogel Phases of PrP C Linked to Conformation Shifts and Triggered by Alzheimer’s Amyloid-β Oligomers. Mol. Cell 72, 426–443 (2018).

26. Matos, C. O. et al. Liquid-liquid phase separation and fibrillation of the prion protein modulated by a high-affinity DNA aptamer. FASEB J. 34, 365–385 (2020).

27. Campana, V., Sarnataro, D. & Zurzolo, C. The highways and byways of prion protein trafficking. Trends Cell Biol. 15, 102–111 (2005).

28. Beutel, O., Maraspini, R., Pombo-García, K., Martin-Lemaitre, C. & Honigmann, A. Phase Separation of Zonula Occludens Proteins Drives Formation of Tight Junctions. Cell (2019) doi:10.1016/j.cell.2019.10.011.

29. do Amaral, M. J., Freire, M. H. O., Almeida, M. S., Pinheiro, A. S. & Cordeiro, Y. Phase separation of the mammalian prion protein: Physiological and pathological perspectives. J. Neurochem. 1–18 (2022) doi:10.1111/jnc.15586.

30. Ivanova, L., Barmada, S., Kummer, T. & Harris, D. A. Mutant Prion Proteins Are Partially Retained in the Endoplasmic Reticulum. J. Biol. Chem. (2001) doi:10.1074/jbc.M106928200.

31. Vendruscolo, M. & Fuxreiter, M. Sequence Determinants of the Aggregation of Proteins Within Condensates Generated by Liquid-liquid Phase Separation: Sequence code of aggregation in protein condensates. J. Mol. Biol. (2022) doi: 10.1016/j.jmb.2021.167201.

32. Bolognesi, B. et al. A concentration-dependent liquid phase separation can cause toxicity upon increased protein expression. Cell Rep. (2016) doi:10.1016/j.celrep.2016.05.076.

33. Paiz, E. A. et al. Beta turn propensity and a model polymer scaling exponent identify intrinsically disordered phase-separating proteins. J. Biol. Chem. (2021) doi:10.1016/j.jbc.2021.101343.

34. Vernon, R. M. C. et al. Pi-Pi contacts are an overlooked protein feature relevant to phase separation. Elife 7, e31486 (2018).

35. Stevens, D. J. et al. Early onset prion disease from octarepeat expansion correlates with copper binding properties. PLoS Pathog. (2009) doi:10.1371/journal.ppat.1000390.

36. Davies, P., Marken, F., Salter, S. & Brown, D. R. Thermodynamic and voltammetric characterization of the metal binding to the prion protein: Insights into pH dependence and redox chemistry. Biochemistry (2009) doi: 10.1021/bi900170n.

37. Sánchez-López, C., Rossetti, G., Quintanar, L. & Carloni, P. Structural determinants of the prion protein N-terminus and its adducts with copper ions. International Journal of Molecular Sciences (2019) doi:10.3390/ijms20010018.

38. Säbel, C. E., Neureuther, J. M. & Siemann, S. A spectrophotometric method for the determination of zinc, copper, and cobalt ions in metalloproteins using Zincon. Anal. Biochem. (2010) doi:10.1016/j.ab.2009.10.037.

39. Xiao, C. Q., Huang, Q., Zhang, Y., Zhang, H. Q. & Lai, L. Binding thermodynamics of divalent metal ions to several biological buffers. Thermochim. Acta (2020) doi:10.1016/j.tca.2020.178721.

40. Viles, J. H. et al. Copper binding to the prion protein: structural implications of four identical cooperative binding sites. Proc. Natl. Acad. Sci. U. S. A. 96, 2042–7 (1999).

41. Sokołowska, M. & Bal, W. Cu(II) complexation by ‘non-coordinating’ N-2-hydroxyethylpiperazine-N’-2-ethanesulfonic acid (HEPES buffer). J. Inorg. Biochem. (2005) doi: 10.1016/j.jinorgbio.2005.05.007.

42. Agarwal, A., Rai, S. K., Avni, A. & Mukhopadhyay, S. An intrinsically disordered pathological prion variant Y145Stop converts into self-seeding amyloids via liquid-liquid phase separation. Proc. Natl. Acad. Sci. U. S. A. 118, e2100968118 (2021).

43. Hochmair, J. et al. Molecular crowding and RNA synergize to promote phase separation, microtubule interaction, and seeding of Tau condensates. EMBO J. (2022) doi:10.15252/embj.2021108882.

44. Hughes, M. P., Goldschmidt, L. & Eisenberg, D. S. Prevalence and species distribution of the low-complexity, amyloid-like, reversible, kinked segment structural motif in amyloid-like fibrils. J. Biol. Chem. (2021) doi:10.1016/j.jbc.2021.101194.

45. Provencher, S. W. & Glöckner, J. Estimation of Globular Protein Secondary Structure from Circular Dichroism. Biochemistry (1981) doi:10.1021/bi00504a006.

46. Nguyen, X. T. A., Tran, T. H., Cojoc, D. & Legname, G. Copper Binding Regulates Cellular Prion Protein Function. Mol. Neurobiol. (2019) doi:10.1007/s12035-019-1510-9.

47. Garnett, A. P. & Viles, J. H. Copper Binding to the Octarepeats of the Prion Protein. J. Biol. Chem. (2003) doi:10.1074/jbc.m209280200.

48. Spiers, J. G., Cortina Chen, H.-J., Barry, T. L., Bourgognon, J.-M. & Steinert, J. R. Redox stress and metal dys-homeostasis appear as hallmarks of early prion disease pathogenesis in mice. Free Radic. Biol. Med. 192, 182–190 (2022).

49. Chan, F. T. S. et al. Protein amyloids develop an intrinsic fluorescence signature during aggregation. Analyst (2013) doi: 10.1039/c3an36798c.

50. Zhang, H. The glassiness of hardening protein droplets. Science (80-.). (2020) doi: 10.1126/science.abe9745.

51. Jawerth, L. et al. Protein condensates as aging Maxwell fluids. Science (80-.). (2020) doi: 10.1126/science.aaw4951.

52. Girelli, A. et al. Microscopic Dynamics of Liquid-Liquid Phase Separation and Domain Coarsening in a Protein Solution Revealed by X-Ray Photon Correlation Spectroscopy. Phys. Rev. Lett. (2021) doi:10.1103/PhysRevLett.126.138004.

53. Moron, M. et al. Gelation Dynamics upon Pressure-Induced Liquid-Liquid Phase Separation in a Water-Lysozyme Solution. J. Phys. Chem. B 126, 4160–4167 (2022).

54. Cipelletti, L. et al. Universal non-diffusive slow dynamics in aging soft matter. Faraday Discuss. (2003) doi:10.1039/b204495a.

55. Ferreira, N. C. et al. A promising antiprion trimethoxychalcone binds to the globular domain of the cellular prion protein and changes its cellular location. Antimicrob. Agents Chemother. 62, 1–17 (2018).

56. Mora, A. K. & Nath, S. SYPRO Orange-a new gold standard amyloid probe. J. Mater. Chem. B (2020) doi:10.1039/d0tb01406k.

57. Maina, M. B. et al. Metal- and UV-Catalyzed Oxidation Results in Trapped Amyloid-β Intermediates Revealing that Self-Assembly Is Required for Aβ-Induced Cytotoxicity. iScience (2020) doi:10.1016/j.isci.2020.101537.

58. McMahon, H. E. M. et al. Cleavage of the amino terminus of the prion protein by reactive oxygen species. J. Biol. Chem. (2001) doi:10.1074/jbc.M007243200.

59. Linsenmeier, L. et al. Diverse functions of the prion protein – Does proteolytic processing hold the key? Biochimica et Biophysica Acta - Molecular Cell Research (2017) doi:10.1016/j.bbamcr.2017.06.022.

60. King, C., Sengupta, P., Seo, A. Y. & Lippincott-Schwartz, J. ER membranes exhibit phase behavior at sites of organelle contact. Proc. Natl. Acad. Sci. U. S. A. (2020) doi:10.1073/pnas.1910854117.

61. Sitar, M. E., Aydin, S. & Çakatay, U. Human serum albumin and its relation with oxidative stress. Clinical Laboratory (2013) doi: 10.7754/Clin.Lab.2012.121115.

62. Bocharova, O. V., Breydo, L., Salnikov, V. V. & Baskakov, I. V. Copper(II) inhibits in vitro conversion of prion protein into amyloid fibrils. Biochemistry 44, 6776–6787 (2005).

63. Thakur, A. K., Srivastava, A. K., Srinivas, V., Chary, K. V. R. & Rao, C. M. Copper alters aggregation behavior of prion protein and induces novel interactions between its N- and C-terminal regions. J. Biol. Chem. 286, 38533–38545 (2011).

64. Tange, H. et al. Liquid liquid phase separation of full-length prion protein initiates conformational conversion in vitro. J. Biol. Chem. 296, 100367 (2021).

65. Passos, Y. M. et al. The interplay between a GC-rich oligonucleotide and copper ions on prion protein conformational and phase transitions. Int. J. Biol. Macromol. 173, 34–43 (2021).

66. Kamps, J. et al. The N-terminal domain of the prion protein is required and sufficient for liquid-liquid phase separation: A crucial role of the Aβ-binding domain. J. Biol. Chem. 297, 100860 (2021).

67. Lin, Y., Fichou, Y., Zeng, Z., Hu, N. Y. & Han, S. Electrostatically Driven Complex Coacervation and Amyloid Aggregation of Tau Are Independent Processes with Overlapping Conditions. ACS Chem. Neurosci. (2020) doi:10.1021/acschemneuro.9b00627.

68. Lipiński, W. P. et al. Biomolecular condensates can both accelerate and suppress aggregation of α-synuclein. Sci. Adv. 8, (2022).

69. Wopfner, F. et al. Analysis of 27 mammalian and 9 avian PrPs reveals high conservation of flexible regions of the prion protein. J.Mol.Biol. 289, 1163–1178 (1999).

70. Polishchuk, E. V. & Polishchuk, R. S. The emerging role of lysosomes in copper homeostasis. Metallomics (2016) doi:10.1039/c6mt00058d.

71. Pauly, P. C. & Harris, D. A. Copper stimulates endocytosis of the prion protein. J. Biol. Chem. (1998) doi:10.1074/jbc.273.50.33107.

72. Petit, C. S. V., Besnier, L., Morel, E., Rousset, M. & Thenet, S. Roles of the cellular prion protein in the regulation of cell-cell junctions and barrier function. Tissue Barriers 1, e24377 (2013).

73. Kouadri, A. et al. Involvement of the Prion Protein in the Protection of the Human Bronchial Epithelial Barrier Against Oxidative Stress. Antioxidants Redox Signal. 31, 59–74 (2019).

74. Mangé, A. et al. Alpha- and beta-cleavages of the amino-terminus of the cellular prion protein. Biol. Cell (2004) doi:10.1016/j.biolcel.2003.11.007.

75. Quaglio, E., Chiesa, R. & Harris, D. A. Copper Converts the Cellular Prion Protein into a Protease-resistant Species That is Distinct from the Scrapie Isoform. J. Biol. Chem. 276, 11432–11438 (2001).

76. Macedo, B., Sant’Anna, R., Navarro, S., Cordeiro, Y. & Ventura, S. Mammalian prion protein (PrP) forms conformationally different amyloid intracellular aggregates in bacteria. Microb. Cell Fact. 14, 174 (2015).

77. Younan, N. D., Nadal, R. C., Davies, P., Brown, D. R. & Viles, J. H. Methionine oxidation perturbs the structural core of the prion protein and suggests a generic misfolding pathway. J. Biol. Chem. (2012) doi:10.1074/jbc.M112.354779.

78. Requena, J. R. et al. Copper-catalyzed oxidation of the recombinant SHa(29-231) prion protein. Proc. Natl. Acad. Sci. U. S. A. (2001) doi:10.1073/pnas.121190898.

79. Kirinčič, S. & Klofutar, C. Viscosity of aqueous solutions of poly(ethylene glycol)s at 298.15 K. Fluid Phase Equilib. (1999) doi:10.1016/S0378-3812(99)00005-9.

80. Legler, D. F. et al. Differential insertion of GPI-anchored GFPs into lipid rafts of live cells. FASEB J. (2005) doi:10.1096/fj.03-1338fje.

81. Schindelin, J. et al. Fiji: An open-source platform for biological-image analysis. Nature Methods (2012) doi:10.1038/nmeth.2019.

