## Extended Data Figures for "Copper drives prion protein phase separation and modulates aggregation"

### Extended Data Fig. 1

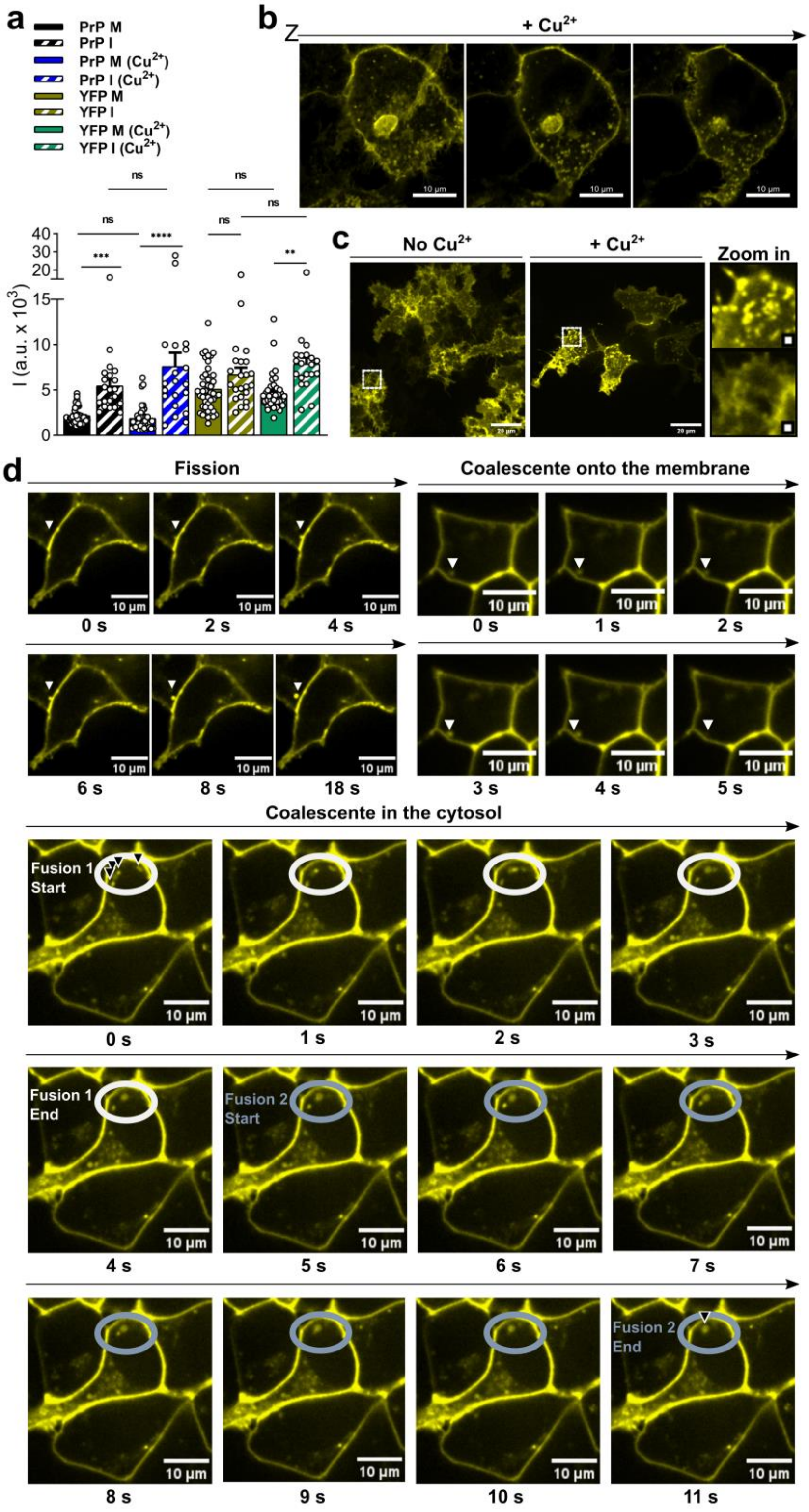


**Extended data Fig. 1 - PrP$^C$-YFP-GPI expression and fission and fusion events**. Relates to Fig. 1. **a,** PrP$^C$-YFP-GPI peak values from line plots analyzed in (**Fig. 1c,d**) show similar membrane and cell-cell interface fluorescence intensities in $CuCl_2$-treated and non-treated cells. ****P<0.0001, ***P<0.001, one-way ANOVA (Tukey). **b** and **c**, Representative Z-stacks of cells incubated for 1 h with 300 µM $CuCl_2$ showing cytosolic PrP$^C$-YFP-GPI punctae and accumulation next to the nucleus (**b**) and the membrane surface plane shows PrP$^C$-YFP-GPI clusters at the cell surface in $CuCl_2$-treated cells (**c**). **d**, Fission and Fusion of cytosolic PrP$^C$-YFP-GPI clusters from/with PM bound PrP$^C$-YFP-GPI. Times indicated below. Cells kept in Opti-MEM in the absence of $Cu^{2+}$.

### Extended Data Fig. 2

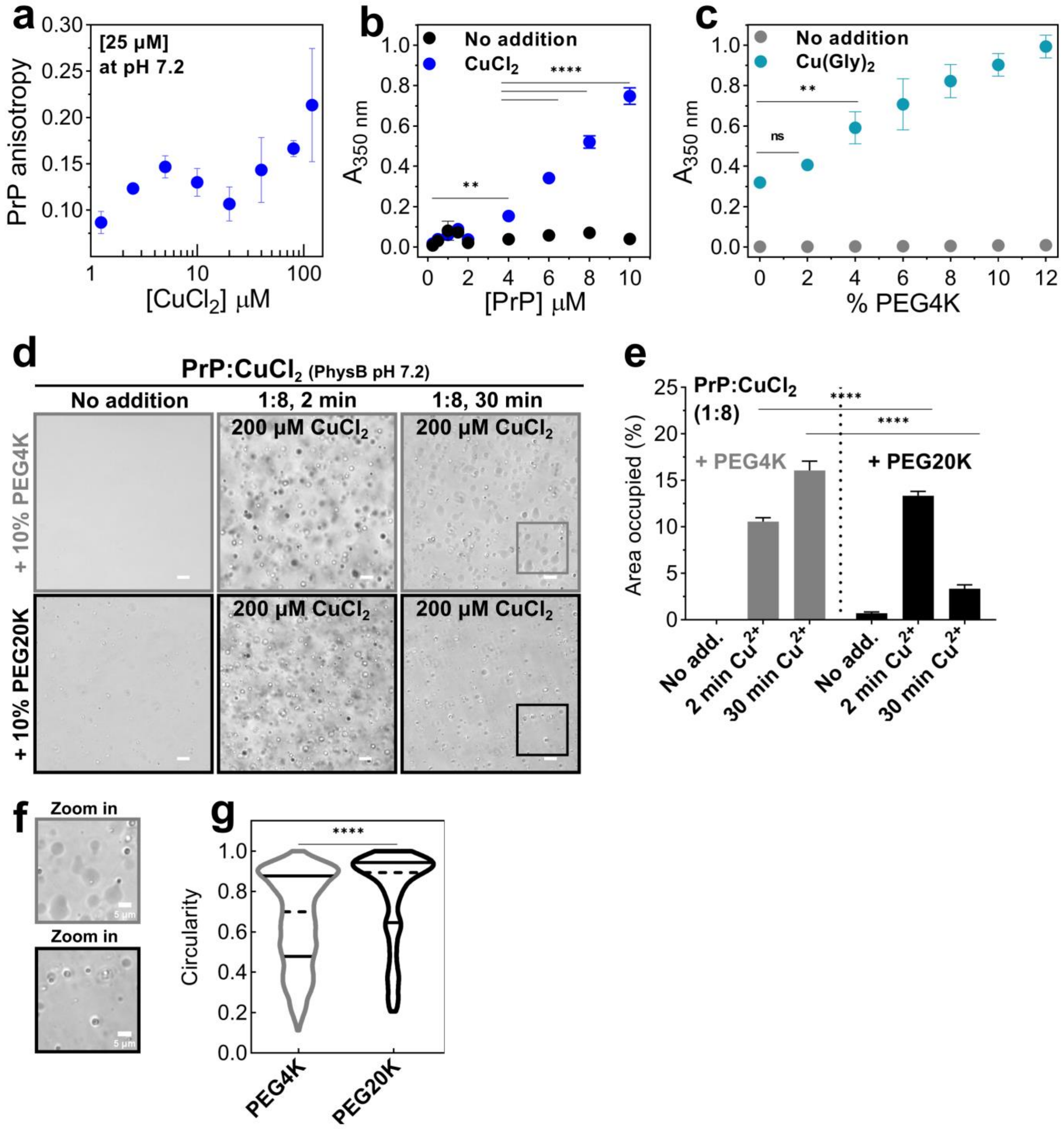


**Extended data Fig. 2 - Effect of solution composition on $Cu^{2+}$-driven PrP condensation**. Relates to Figure 2. **a,** Tryptophan anisotropy (excitation 280 nm; emission 353 nm) of 25 µM PrP upon $CuCl_2$ titration. **b,** Turbidity (absorbance at 350 nm) measured after 30 minutes sample preparation with increasing PrP concentrations in PhysB pH 7.2 (black) and with 80 µM $CuCl_2$ addition (blue). **c,** Turbidity of 25 µM PrP with 200 µM $Cu(Gly)_2$ (1:8 in 20 mM HEPES pH 7.2) as a function of PEG4K concentration (w/v). **d**, Phase contrast images of 25 µM PrP with 200 µM $CuCl_2$ in PhysB containing 10% (w/v) PEG4K or PEG20K. **e,** Quantification of surface area occupied by condensates. Data shown as mean ± SD, n= 6 images per condition. **f,** Zoomed regions from indicated ROIs in (**d**) show wetting of the condensates on the coverslip. **g,** Violin plot of circularity distribution from surface attached condensates shown in (**d**) (third column, 30 min). Solid lines indicate quartiles, dashed lines show medians. n= 927 condensates for PEG4K, and 357 condensates for PEG20K from 6 different images each. Data analyzed by one-way ANOVA (Tukey) in (**b**, **c**, **e**) and two-tailed Student's t (**g**). **P<0.001, ****P < 0.0001, ns, not significant.

### Extended Data Fig. 3

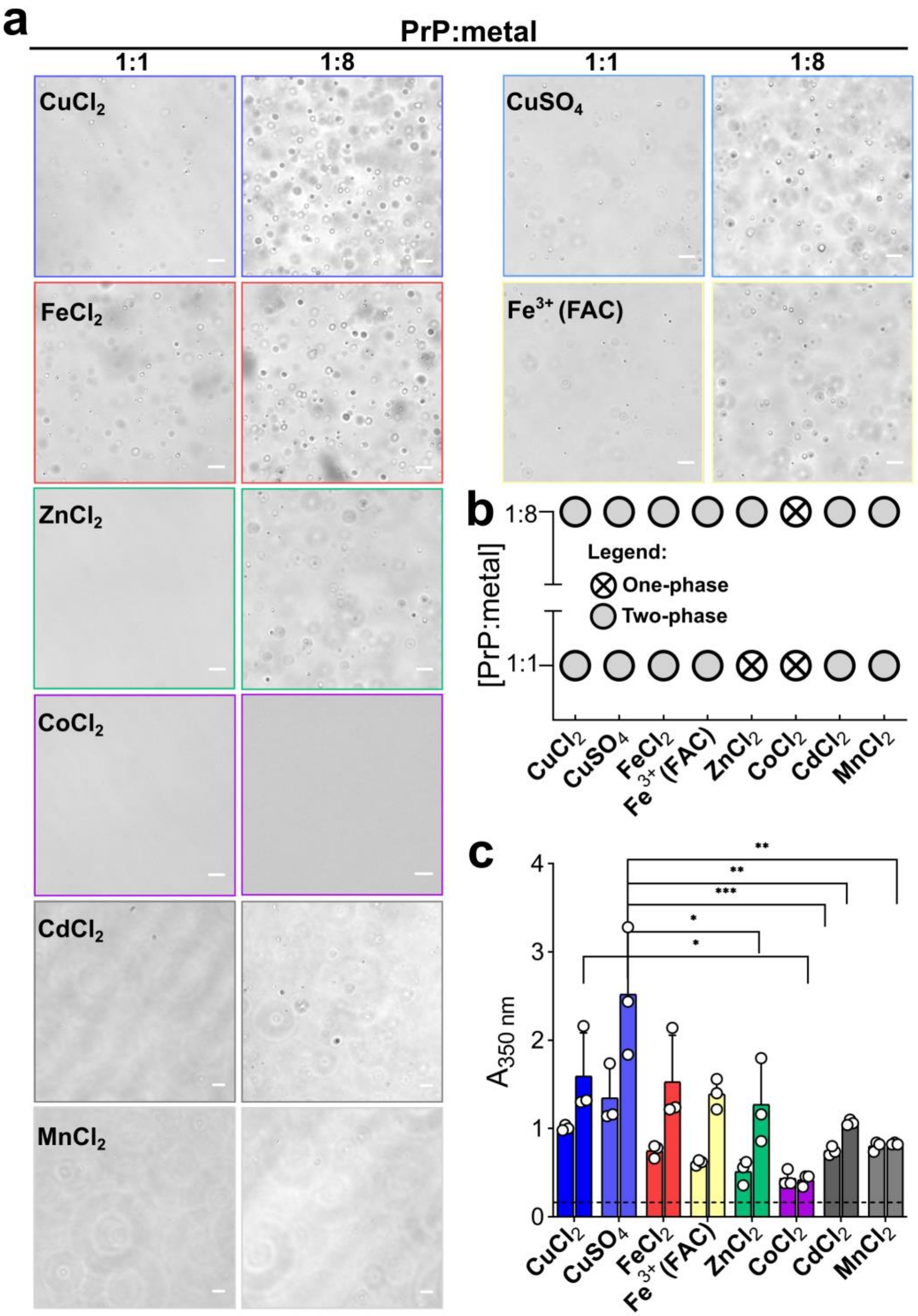


**Extended data Fig. 3 - Pathophysiological relevant divalent metals trigger PrP condensation. a,** Representative phase contrast images are shown for 25 µM PrP with indicated metals at 25 (1:1) or 200 µM (1:8). Scale bars, 10 µm. **b,** Diagram based in 'a' (n=3 independent experiments). **c**, Turbidity (absorbance at 350 nm) with specified metals at 1:1 (first bar) and 1:8 (second bar) molar ratios. Data shown as mean ± SD, (n=3), one-way ANOVA (Tukey). Significant comparisons are shown for the 1:8 molar ratio (***$P < 0.001$; **$P < 0.01$; *$P < 0.05$). For the 1:1 molar ratio, results are as follow: *$P < 0.05$ ($CuCl_2$ vs. $CoCl_2$; $ZnCl_2$ vs. $CuSO_4$), **$P < 0.01$ ($CuSO_4$ vs. $CdCl_2$; $CuSO_4$ vs. $MnCl_2$), ***$P < 0.001$ ($CuSO_4$ vs. $CoCl_2$). Non significant (ns) comparisons for both molar ratios ($CuCl_2$ vs. $CuSO_4$; $CuCl_2$ vs. $Fe^{3+}$(FAC); $CuCl_2$ vs. $FeCl_2$; $CuCl_2$ vs. $CdCl_2$; $CuCl_2$ vs. $MnCl_2$; $ZnCl_2$ vs. $CoCl_2$; $ZnCl_2$ vs. $Fe^{3+}$(FAC); $ZnCl_2$ vs. $FeCl_2$; $ZnCl_2$ vs. $CdCl_2$; $ZnCl_2$ vs. $MnCl_2$; $CoCl_2$ vs. $Fe^{3+}$(FAC); $CoCl_2$ vs. $FeCl_2$; $CoCl_2$ vs. $CdCl_2$; $CoCl_2$ vs. $MnCl_2$; $Fe^{3+}$(FAC) vs. $FeCl_2$; $Fe^{3+}$(FAC) vs. $CdCl_2$; $Fe^{3+}$(FAC) vs. $MnCl_2$; $FeCl_2$ vs. $CdCl_2$; $FeCl_2$ vs. $MnCl_2$; $CdCl_2$ vs. $MnCl_2$) together with ($CuCl_2$ vs. $ZnCl_{2;}$; $CuSO_4$ vs. $Fe^{3+}$(FAC) and $CuSO_4$ vs. $FeCl_2$) for the 1:8 molar ratio. Samples were prepared in PhysB, pH 7.2 at 25 °C for 1 h before imaging or turbidimetry.

### Extended Data Fig. 4

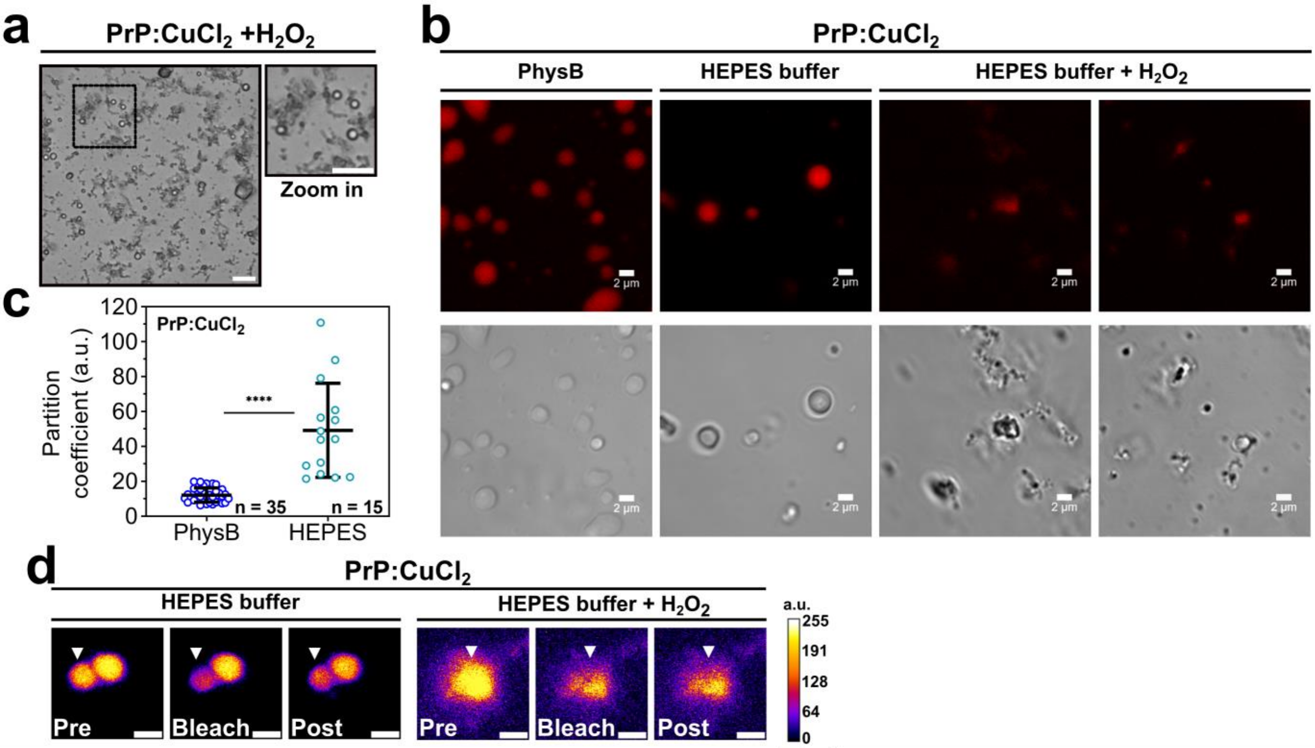


**Extended data Fig. 4 - Additional microscopy images for PrP:$Cu^{2+}$ with $H_2O_2$ and FRAP images. Relates to Figure 4. a,** Phase contrast of 25 µM PrP with 200 µM $CuCl_2$ and 10 mM $H_2O_2$ in PhysB pH 7.2 showing condensates and aggregates after 30 minutes preparation. Scale bars, 10 µm. **b,** Representative Fluorescence and DIC images of 10 µM PrP with 80 µM $CuCl_2$ in the indicated buffers containing or not 10 mM $H_2O_2$. Scale bars, 2 µm. **c,** Partition coefficients obtained from images as shown in 'b', with exception of the sample with addition of $H_2O_2$ due to the circularity (> 0.5) and size (> 1.0 µm$^2$) considered in this analysis; n = 35 condensates for physiological-like buffer pH 7.2; n = 15 for 20 mM HEPES buffer pH 7.4; ****P < 0.001 (unpaired t-test). **d**, Representative images from FRAP experiment (refers to Figure 4e) before bleaching (pre), just after bleaching (bleach) and after recovery (t= 50 s; post). Arrows indicate circular ROIs (diameter 0.8 µm) selected for FRAP. Scale bars, 2 µm. All samples were prepared at 25 °C.

### Extended Data Fig. 5

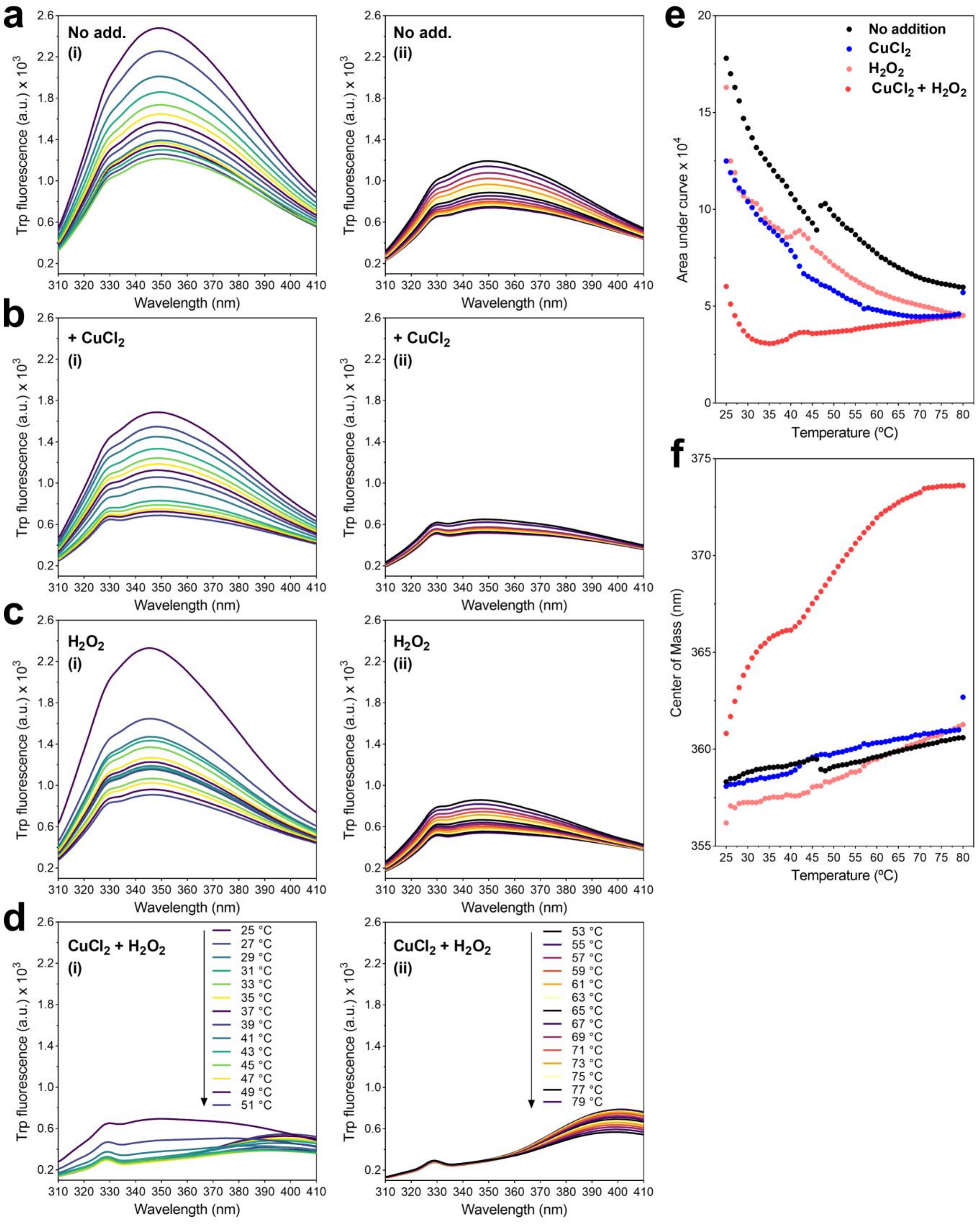


**Extended data Fig. 5 - $Cu^{2+}$ and $H_2O_2$ immediately induce changes in intrinsic tryptophan fluorescence evidenced by dityrosine formation**. Intrinsic fluorescence of tryptophan (excitation at 295 nm) upon slowly heating PrP samples at 2.5 μM in steps of 1°C. For simplicity, spectra are shown in steps of 2 °C; 25-51 °C (first row, 'i') and 53-79 °C (second row, 'ii'). **a**, PrP in PhysB pH 7.2. **b**, PrP:$CuCl_2$ (1:8 molar ratio). **c**, PrP upon addition of 10 mM $H_2O_2$**. d,** PrP:$CuCl_2$ (1:8) + 10 mM $H_2O_2$. **e**, Area under curve of tryptophan fluorescence emission *versus* temperature extracted from 'a' to 'd'. **f**, Center of mass *versus* temperature extracted from 'a' to 'd'.
