## Supplementary Material for "Copper drives prion protein phase separation and modulates aggregation"

Supplementary Figures 1 to 7

Supplementary Table 1

Supplementary Videos 1 to 4

References

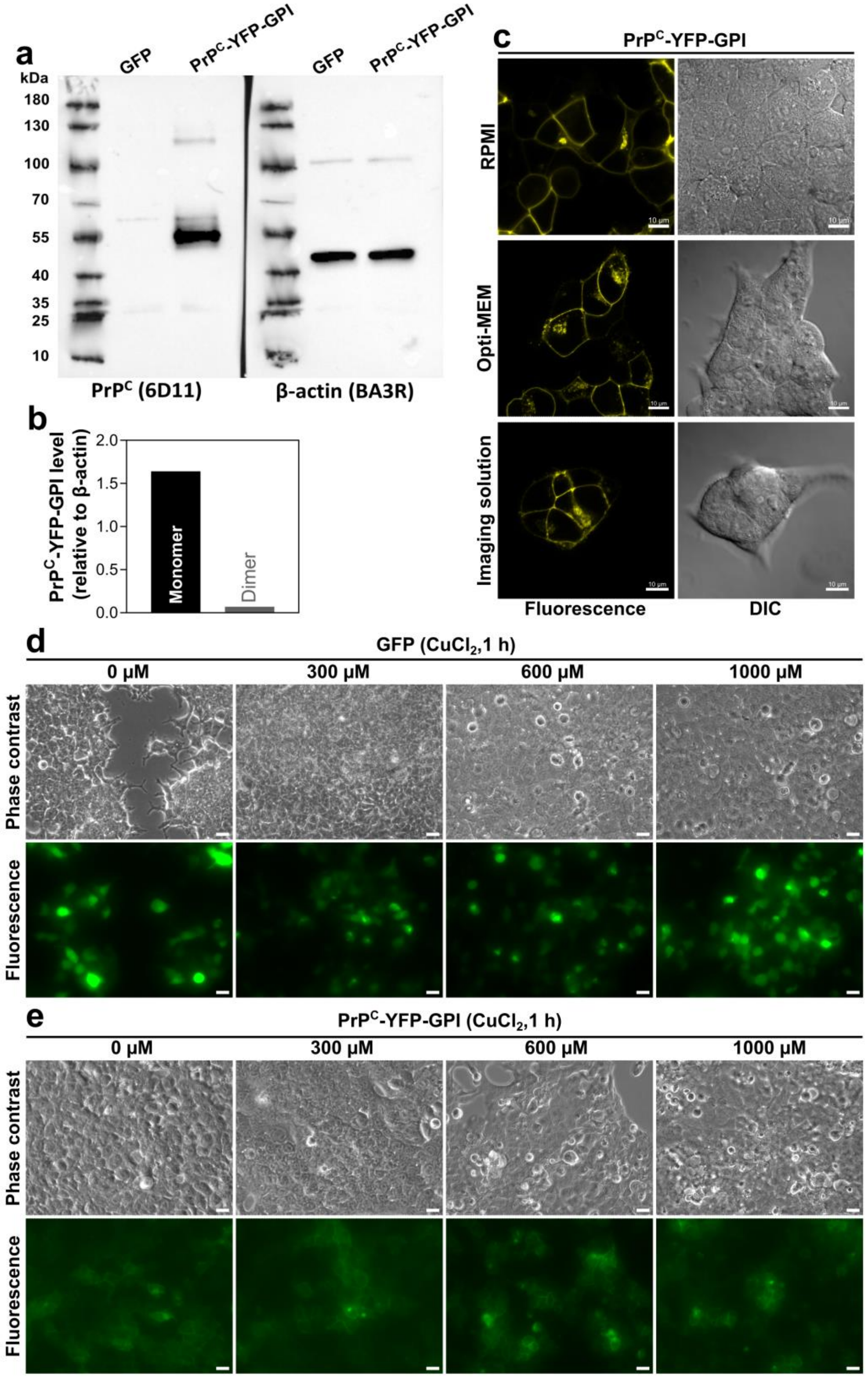


**Supplementary Fig. 1 - Comparison of $CuCl_2$ treatment in GFP versus PrP$^{C}$-YFP-GPI expressing HEK293 cells.** Relates to Fig. 1**. a**, Western blot of cell lysates from HEK293 cells expressing GFP or PrP$^{C}$-YFP-GPI using anti-PrP$^{C}$ (6D11) and anti-β-actin (BA3R) as loading control. **b**, Level of PrP$^{C}$-YFP-GPI monomers and dimers relative to β-actin analyzed from (a). **c**, confocal imaging of live HEK293 cells expressing PrP$^{C}$-YFP-GPI (for 48 h) maintained in RPMI (top) or Opti-MEM (middle) or Imaging solution (bottom) for 2 h. **e**, Representative images collected in parallel with cell viability assay in **Fig. 1g**. Scale bars, 10 μm.

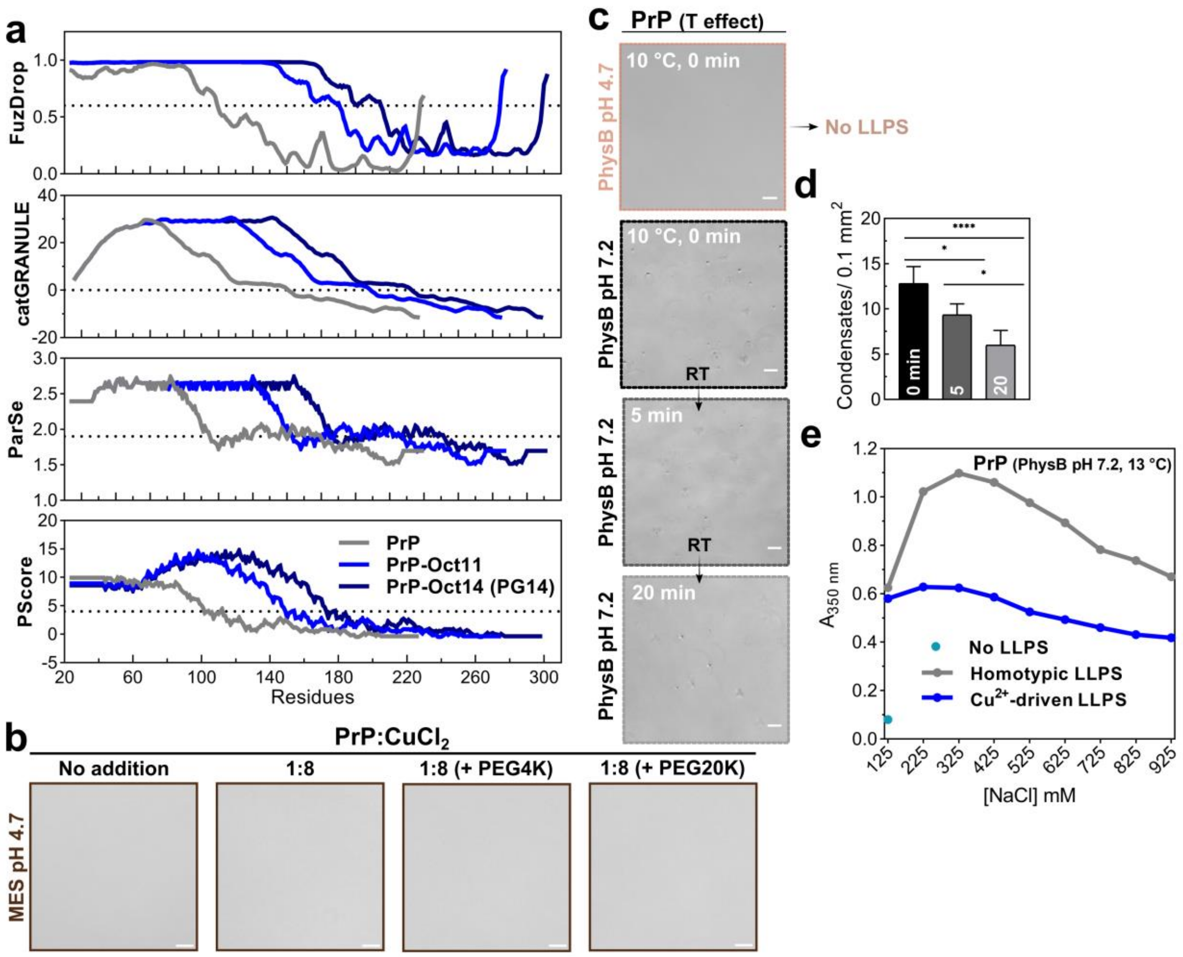


**Supplementary Fig. 2 - LLPS predictions for PrP mutants and in vitro PrP condensation triggered by low temperature and salt**. Relates to Fig. 2a. **a**, LLPS propensities prediction for wild-type PrP (PrP; residues 23-231), mutants PrP-Oct11 (6 extra octarepeats) and PrP-Oct14 (9 extra octarepeats; PG14) by FuzDrop (Vendruscolo *et al*., 2022), catGRANULE (Bolognesi *et al*., 2016), ParSe (Paiz *et al*., 2021) and PScore (Vernon *et al*., 2018). Residues with scores above indicated dotted lines are prone to mediate LLPS. **b**, Relates to Figure 2i. 25 µM PrP with 200 µM $CuCl_2$ in 25 mM MES pH 4.7 (without salt) containing no, or 10% PEG4K or PEG20K. **c** to **e,** Low temperature promotes PrP homotypic LLPS in PhysB pH 7.2. **c,** Phase contrast images of 25 µM PrP incubated for 1 h at 0 °C then imaged at 25 °C over time in PhysB (pH 7.2, and pH 4.7 in which sodium phosphate was replaced to sodium acetate). Temperature (T) of first sample was 10 °C. **d,** Condensate's quantification as a function of time from (c). RT, room temperature. Data shown as mean ± SD (n= 3 images), one-way ANOVA with Tukey test. ****$P < 0.0001$, *$P < 0.05$. **e**, Ionic strength effect in homotypic (PrP) vs. heterotypic (PrP:$Cu^{2+}$) condensation. Turbidity of PrP (71 µM; grey) and PrP:$CuCl_2$ (1:1; 25 µM each; blue) with NaCl titration (indicated in x-axis). To obtain the same degree of homotypic and heterotypic LLPS, PrP concentration was adjusted to have the same turbidity at the starting point. The turquoise circle (on y-axis) corresponds to the turbidity of a PrP only solution at 25 µM (for comparison to the blue curve). PhysB had KCl replaced by NaCl. Samples were pre-equilibrated at 13 °C (1 hour).

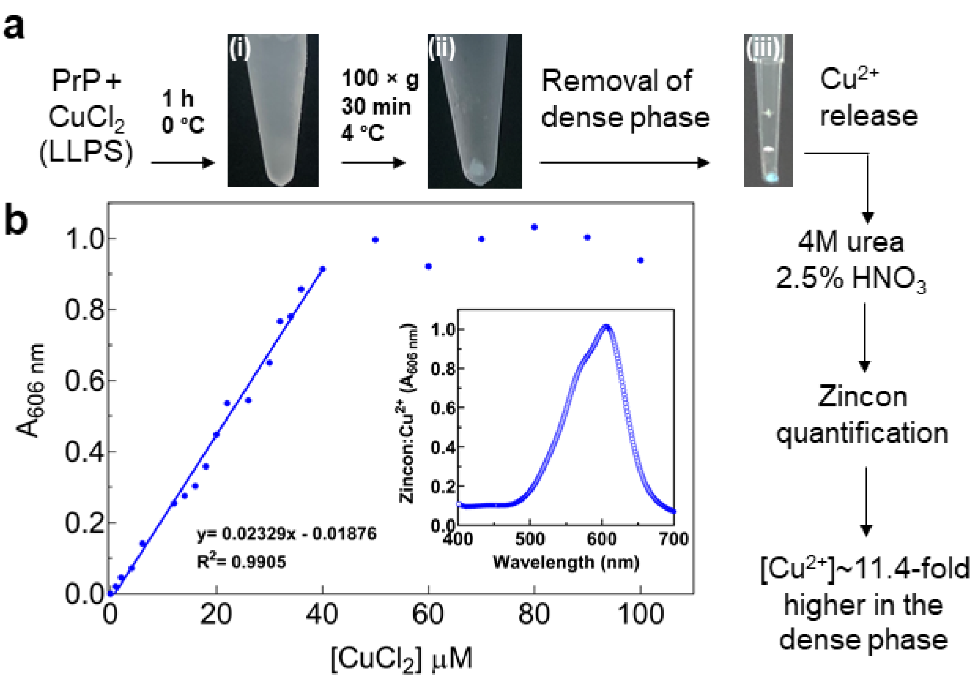


**Supplementary Fig. 3 - $Cu^{2+}$ is concentrated in the PrP condensates.** Relates to Fig. 2e,f. **a,** Workflow of $Cu^{+2}$ quantification in the dense phase by Zincon. 25 µM PrP with 200 µM $CuCl_2$ in 10 mM HEPES pH 7.4, 120 mM KCl, 5 mM NaCl, 6% (w/v) PEG4000 was incubated in an ice bath for 1 h (cloudy solution, photo 'i') followed by centrifugation (note the blue pellet, photo 'ii') and removal of the dense phase (blue pellet in the tip bottom, photo 'iii') which was weighted and incubated with urea and $HNO_3$ to release bound $Cu^{2+}$ followed by Zincon quantification. **b**, Calibration curve of Zincon with $CuCl_2$. in 50 mM borate buffer pH 9 containing 8 M urea. Reactions were prepared with 40 µM Zincon with increasing $CuCl_2$ concentrations for 30 min prior to absorbance readings at 606 nm ($\lambda_{max}$ ). Inset: Zincon (40 µM) absorbance spectra with excess of $CuCl_2$ (200 µM). The maximum at 606 nm was used for quantification.

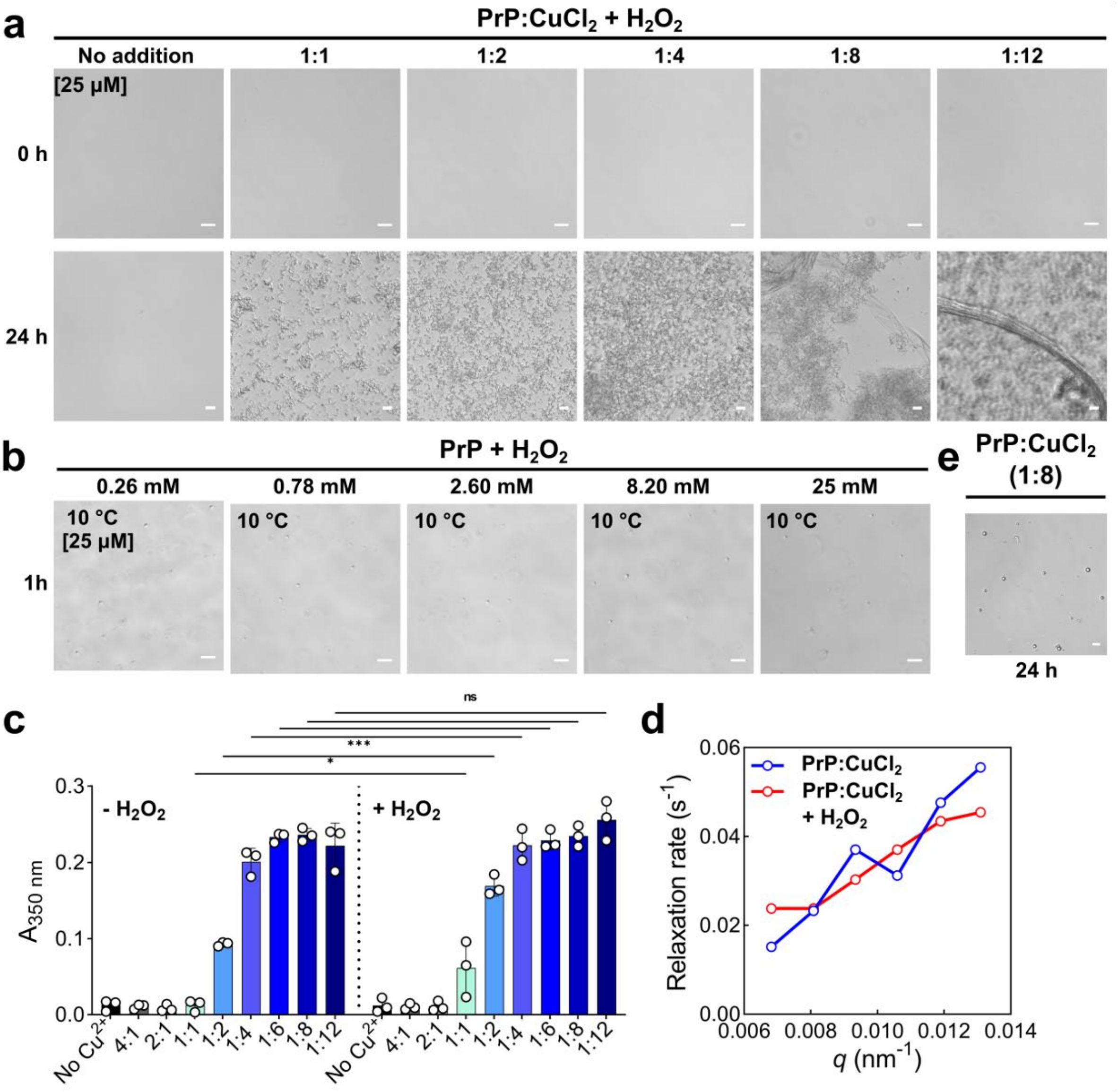


**Supplementary Fig. 4 - Hydrogen peroxide effect on PrP:$Cu^{2+}$ LLPS.** Relates to Fig. 4. **a**, Additional time points with the same samples from **Fig. 4a** of 25 µM PrP with 25-300 µM $CuCl_2$ (molar ratios indicated above) in the presence of 10 mM $H_2O_2$. **b**, Phase contrast images of 25µM PrP with 0.26 to 25 mM $H_2O_2$ incubated for 1 h at 0°C, then imaged after 5 minutes when the temperature was ~10°C. Few condensates are formed at low temperatures and $H_2O_2$ did not modified LLPS. Scale bars, 10 µm. **c**, Same PrP:$Cu^{2+}$ turbidity data shown in **Fig. 2c** and **Fig. 4b** for statistical comparison. PrP:$CuCl_2$ molar ratios are shown on the x-axis. Data analyzed by one-way ANOVA with Tukey test for multiple comparison. *P < 0.05; ***P < 0.001; ns, not significant. **d**, XPCS relaxation dynamics of PrP:$CuCl_2$ with and without $H_2O_2$. The graph shows the behavior of relaxation rate over the whole *q* range in which the dynamic is accessible. **e**, Phase contrast image of 25 µM PrP with 200 µM $CuCl_2$ (1:8) acquired after 24h, PrP:$Cu^{2+}$ droplets in the bottom surface remain spherical. Scale bars, 10 µm. All samples prepared in PhysB pH 7.2 at 25 °C, unless otherwise stated.

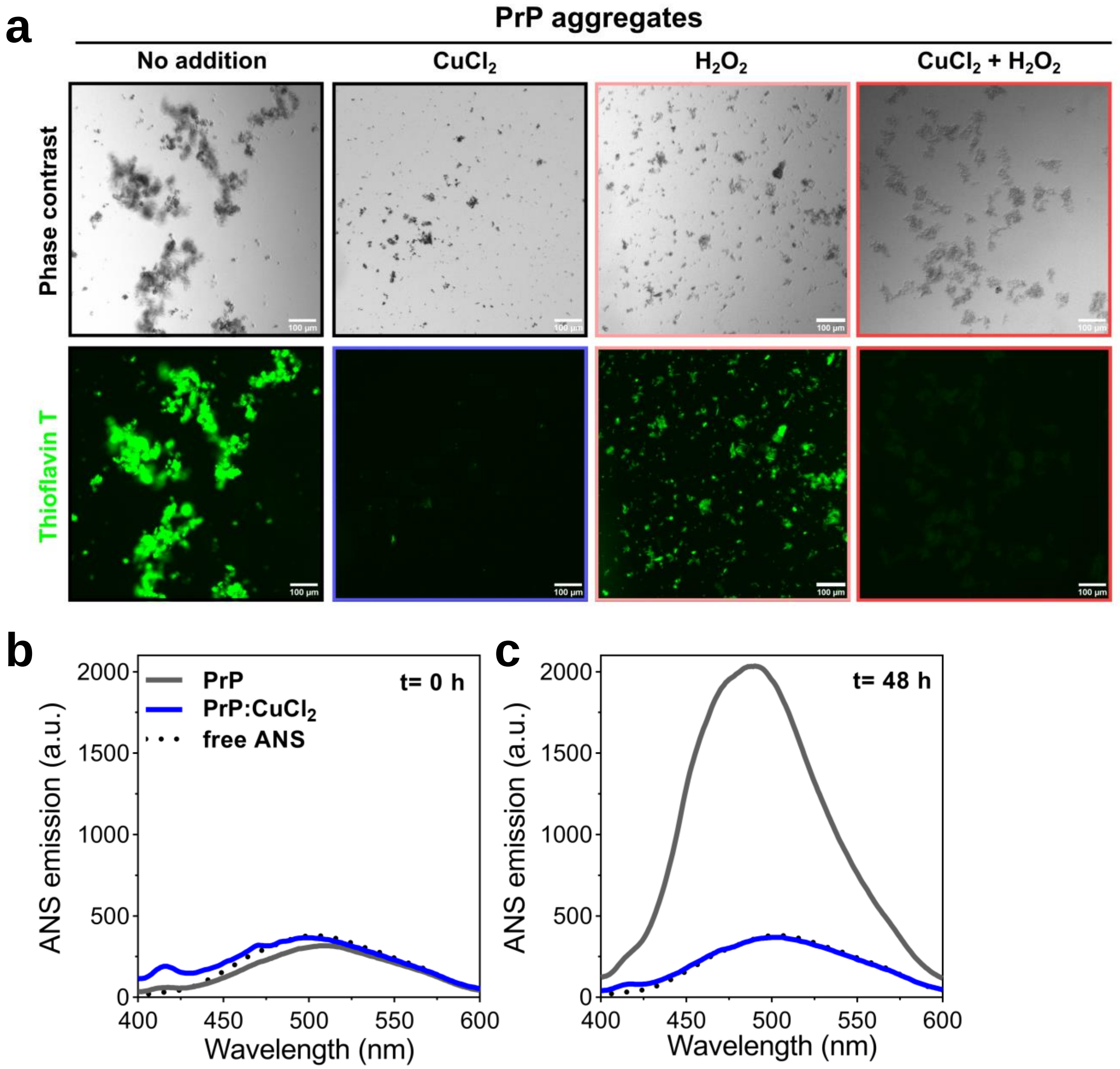


**Supplementary Fig. 5 - PrP aggregation in the presence of $Cu^{2+}$ does not lead ThT-positive and partially unfolded aggregates.** Relates to Fig. 5. Soluble PrP was incubated with 0.1% seed of pre-aggregated recombinant PrP (produced as in Ferreira *et al*., 2018) under continuous agitation at 42ºC for 48 h in the presence or absence of $CuCl_2$ (1:8 molar ratio), 10 mM $H_2O_2$ or both as illustrated in Fig. 5a. **a**, Representative phase contrast images and corresponding extrinsic fluorescence with 50 µM thioflavin T. **b** and **c**, Fluorescence emission spectra (excitation 360 nm) of 30 µM 8-anilinonaphthalene-1-sulfonic acid (ANS) in 20 µM PrP, immediately (**b**) or 48 h (**c**) after the aggregation protocol. Emission of free ANS in buffer (dashed line). All samples and spectra acquisition in PhysB pH 7.2 at 25ºC .

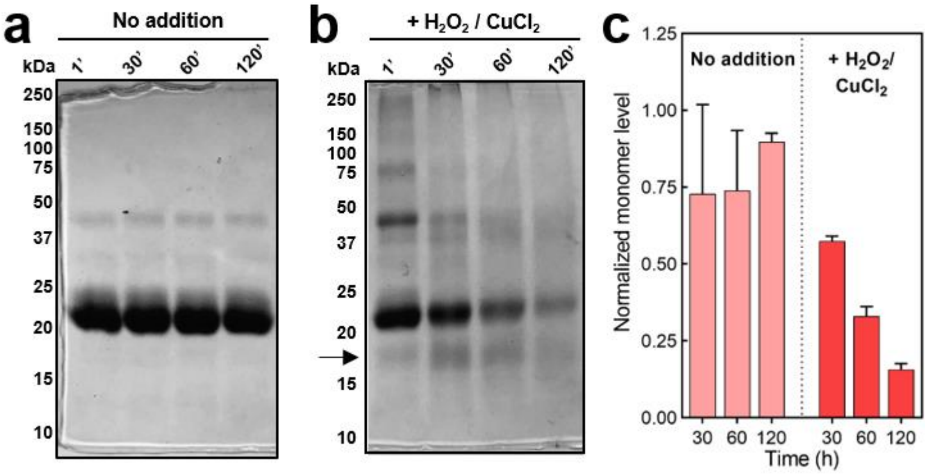


**Supplementary Fig. 6 - $H_2O_2$ and $Cu^{2+}$ promote recombinant PrP cleavage. a** and **b,** 15% SDS-PAGEs stained with colloidal Coomassie Blue G-250. Time-course of 25 µM PrP solutions and upon addition of 25 µM $CuCl_2$ (1:1 molar ratio) and 10 mM $H_2O_2$. Each lane was loaded with 11.5 µg protein. **c,** Quantification of PrP monomer band using samples ran in SDS-PAGEs ('a' and 'b' and other gels not shown). Values normalized to the first time point (1 minute). Data is shown as mean ± SEM (n=2 independent experiments). All samples were prepared in PhysB pH 7.2 in which KCl was replaced to NaCl to avoid SDS precipitation. Samples incubated at 25 °C and retrieved at the specified time points followed by methanol precipitation. Position of molecular weight standards (in kDa) are indicated to the left of the gels.

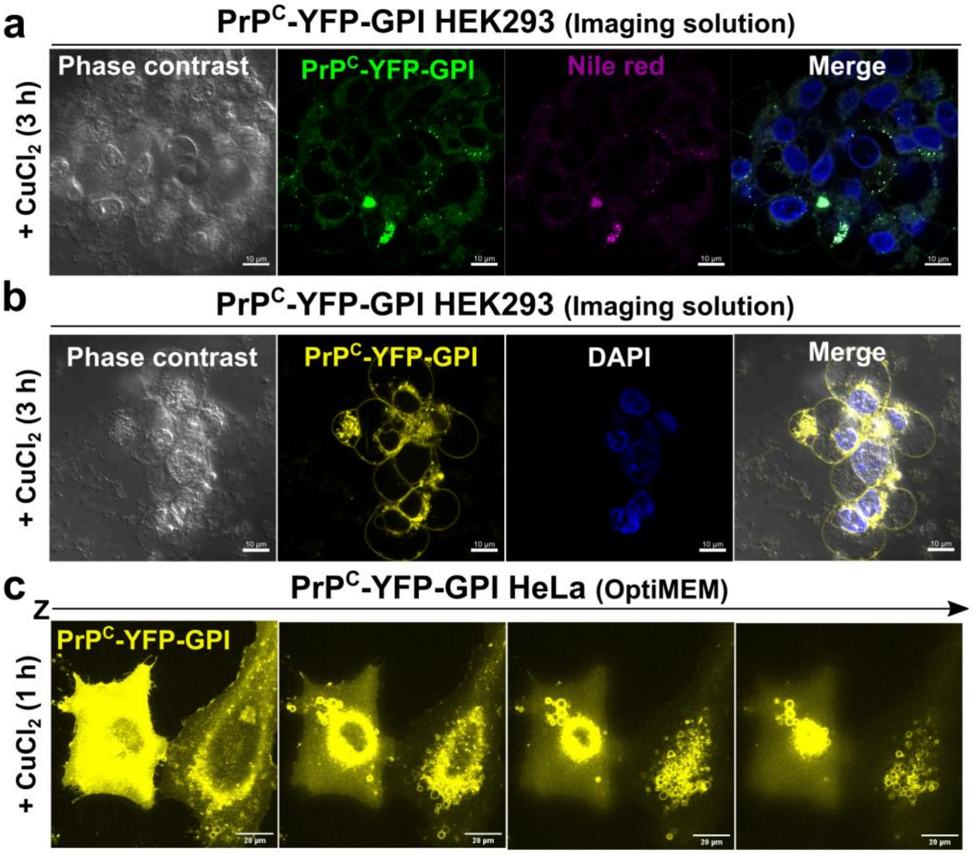


**Supplementary Fig. 7 - Phenotypic findings of PrP$^{C}$-YFP-GPI cells with $Cu^{2+}$ incubation**. Relates to Fig. 6. Live cell imaging of cells transfected with PrP$^{C}$-YFP-GPI (48 h p.t.). **a**, 3 h $Cu^{2+}$ treatment in imaging solution followed by addition of Nile Red. **b**, 3 h $Cu^{2+}$ treatment in imaging solution of HEK293 cells showing PrPC-YFP-GPI aggregates around the nucleus. **c**, Optical sections obtained by confocal laser scanning of HeLa cells incubated with 300 µM $CuCl_2$ for 1 h .

**Supplementary Table 1. Composition of the physiological buffer (PhysB) used in this study and relevant physiological concentrations of copper ions reported in the literature.**

| **Physiological buffer (PhysB)** | |
|---|---|
| **Salt** | Concentration |
| **HEPES pH 7.2*** | 10 mM |
| **KCl** | 120 mM |
| **NaCl** | 5 mM |
| **$CaCl_2$** | 150 µM |
| **$MgCl_2$** | 5 mM |
| **PEG4K** | 60 mg/mL (= 6% w/v) |
| **Copper ions** | |
| **Location** | Concentration |
| **CSF** | ~70 µM (Kardos *et al*. 1989) |
| **Human brain** | ~80 µM (Stöckel *et al*. 1998);<br>3-5 µg/g wet tissue (Kardos *et al*. 2018) |
| **Locus coeruleus** | ~860 ng/mg of wet tissue<br>(Zecca *et al*. 2004) |
| **Substantia nigra** | 1200 ng/mg of wet tissue<br>(Zecca *et al*. 2004) |
| **Synaptic cleft** | 100-250 µM (Kardos *et al*. 1989)<br>200-400 µM (Hopt *et al*. 2003) |
| **Vesicle** | ~291 µM (Hopt *et al*. 2003) |
| **Synaptosome** | ~48 µM (Hopt *et al*. 2003)<br>~15 µM (Herms *et al*. 1999) |

*Replaced to $K_2HPO_4/KH_2PO_4$ in some experiments due to technical hindrance. Buffer composition was adapted from the 'cytomix' buffer (Knight and Scrutton, 1986; Van Den Hoff *et al*., 1992) together with the findings on the ionic composition and the pH values reported for mammalian cells and reviewed in (Theillet *et al*., 2014).

**Supplementary Video 1 –** Time lapse of HEK293 cells expressing $PrP^{C}$-YFP-GPI and treated with 300 µM $CuCl_2$ (for 1 h) showing accumulation in cell-cell interfaces and interplay between condensates and the enriched $PrP^{C}$ species at the plasma membrane. Scale bar, 10 µm. Time stamp indicated on top left.

**Supplementary Video 2 –** An isolated pair of HEK293 cells expressing $PrP^{C}$-YFP-GPI demonstrate the clear enrichment of $PrP^{C}$-YFP-GPI at the cell-cell interface upon $Cu^{2+}$ treatment (300 µM $CuCl_2$ for 1 h) and the quickly $PrP^{C}$ condensates membrane-cytosol exchange. Scale bar, 10 µm. Time stamp indicated on top left.

**Supplementary Video 3 –** PrP:$Cu^{2+}$ condensates wet a poly-D-lysine coated glass surface. 20 µM PrP (spiked with 0.1% PrP-Alexa647) with 160 µM $CuCl_2$ (1:8 molar ratio) in 10 mM HEPES, pH 7.2, no salt. Time lapse started after 15 min sample preparation at 25 ºC. Time stamp indicated on top left.

**Supplementary Video 4 –** HEK293 cells expressing $PrP^{C}$-YFP-GPI upon 3 h $Cu^{2+}$ treatment (300 µM $CuCl_2$) show hollow structures in the cytosol (indicated by the ROIs).

### References

Bolognesi, B., Gotor, N. L., Dhar, R., Cirillo, D., Baldrighi, M., Tartaglia, G. G., & Lehner, B. (2016). A concentration-dependent liquid phase separation can cause toxicity upon increased protein expression. *Cell reports*, *16*(1), 222-231.

Ferreira, N. C., Ascari, L. M., Hughson, A. G., Cavalheiro, G. R., Góes, C. F., Fernandes, P. N., ... & Cordeiro, Y. (2018). A promising antiprion trimethoxychalcone binds to the globular domain of the cellular prion protein and changes its cellular location. *Antimicrobial agents and chemotherapy*, *62*(2), e01441-17.

Herms, J., Tings, T., Gall, S., Madlung, A., Giese, A., Siebert, H., et al. (1999). Evidence of presynaptic location and function of the prion protein. *J. Neurosci.* doi:10.1523/jneurosci.19-20-08866.1999.

Hopt, A., Korte, S., Fink, H., Panne, U., Niessner, R., Jahn, R., et al. (2003). Methods for studying synaptosomal copper release. *J. Neurosci. Methods*. doi:10.1016/S0165-0270(03)00173-0.

Kardos, J., Héja, L., Simon, Á., Jablonkai, I., Kovács, R., and Jemnitz, K. (2018). Copper signalling: causes and consequences. *Cell Communication and Signaling*, 16, 1-22.

Kardos, J., Kovács, I., Hajós, F., Kálmán, M., and Simonyi, M. (1989). Nerve endings from rat brain tissue release copper upon depolarization. A possible role in regulating neuronal excitability. *Neurosci. Lett.* doi:10.1016/0304-3940(89)90565-X.

Knight, D. E., and Scrutton, M. C. (1986). Gaining access to the cytosol: The technique and some applications of electropermeabilization. *Biochem. J.* doi:10.1042/bj2340497.

Paiz, E. A., Allen, J. H., Correia, J. J., Fitzkee, N. C., Hough, L. E., & Whitten, S. T. (2021). Beta turn propensity and a model polymer scaling exponent identify intrinsically disordered phase-separating proteins. *Journal of Biological Chemistry*, *297*(5).

Stöckel, J., Safar, J., Wallace, A. C., Cohen, F. E., and Prusiner, S. B. (1998). Prion protein selectively binds copper(II) ions. *Biochemistry*. doi:10.1021/bi972827k.

Van Den Hoff, M. J. b., Moorman, A. F. M., and Lamers, W. H. (1992). Electroporation in "intracellular" buffer increases cell survival. *Nucleic Acids Res.* doi:10.1093/nar/20.11.2902.

Vendruscolo, M., & Fuxreiter, M. (2022). Sequence determinants of the aggregation of proteins within condensates generated by liquid-liquid phase separation. *Journal of Molecular Biology*, *434*(1), 167201.

Vernon, R. M., Chong, P. A., Tsang, B., Kim, T. H., Bah, A., Farber, P., ... & Forman-Kay, J. D. (2018). Pi-Pi contacts are an overlooked protein feature relevant to phase separation. *elife*, *7*.

Theillet, F. X., Binolfi, A., Frembgen-Kesner, T., Hingorani, K., Sarkar, M., Kyne, C., et al. (2014). Physicochemical properties of cells and their effects on intrinsically disordered proteins (IDPs). *Chem. Rev.* 114, 6661–6714. doi:10.1021/cr400695p.

Zecca, L., Stroppolo, A., Gatti, A., Tampellini, D., Toscani, M., Gallorini, M., et al. (2004). The role of iron and molecules in the neuronal vulnerability of locus coeruleus and substantia nigra during aging. *Proc. Natl. Acad. Sci. U. S. A.* doi:10.1073/pnas.0403495101.
